# A genome wide antigen-antibody screen identifies a yeast-based therapeutic vaccine candidate for Chagas disease

**DOI:** 10.64898/2026.06.12.731947

**Authors:** Mira Loock, Lissa Cruz-Saavedra, Valeria Bernal Araujo, Igor Cestari

## Abstract

Chagas disease is caused by *Trypanosoma cruzi* infection and results in decades-long, chronic, debilitating, and lethal disease. There is no vaccine available, largely due to the lack of vaccine targets. We generated a *T. cruzi* genome-wide library for yeast surface display and screened it with antibodies from Chagas disease patients to identify antigen-antibody interactions. We identified hundreds of *T. cruzi* antigens that are immunogenic in humans and mapped their antibody-binding sites at nucleotide resolution. Hundreds of immunogens were conserved across strains, divergent from human proteins, and expressed across all *T. cruzi* infectious forms, revealing potential vaccine targets. We vaccinated mice with a 63-amino-acid-long immunogenic region using recombinant protein (rIR1) or yeast surface expression (yIR1). yIR1 surpassed rIR1, reducing parasitemia in blood and muscle tissues in a prophylactic, acute-stage vaccination model, validating IR1 as an immunogen and yeast as a vaccine vehicle for Chagas disease. Notably, therapeutic vaccination of chronically infected mice with yIR1 cleared parasites from muscle tissues and was accompanied by the production of α-IR1 antibodies and an increased proportion of CD8^+^ T cells, indicating that a vaccine for treating chronic Chagas disease may be feasible. The antigen-antibody screen revealed hundreds of immunogens for vaccine, diagnostic, and biomarker discovery, and uncovered yIR1 as a therapeutic vaccine candidate for Chagas disease.

**One Sentence Summary:** A yeast-based therapeutic vaccine candidate for treating chronic Chagas disease

## INTRODUCTION

Chagas disease (CD) is caused by infection with the single-celled protozoan parasite *Trypanosoma cruzi*. CD affects ∼10.5 million people, largely in South and Central America (*1, 2*). It has spread to the United States of America, where vectorial transmission has been detected in the past decade (*2, 3*), as well as to Canada, Europe, Asia, and Australia (*1*), becoming a global health concern with an economic burden estimated at over 7 billion annually (*4, 5*). *T. cruzi* is transmitted through the bite of a triatomine vector, or by consuming contaminated food or drinks, through blood transfusion, or congenitally (*1*). Upon infection, patients enter an acute stage characterized by flu-like symptoms that are often undiagnosed, which then progresses to a decades-long chronic stage with asymptomatic (∼60%) or symptomatic (∼40%) forms. Chronic CD patients typically develop cardiac diseases, e.g., cardiomegaly leading to heart failure, or gastrointestinal disorders, such as megaesophagus, megacolon, and enteropathy (*1*).

Benznidazole and nifurtimox are the available drugs for treating CD, with benznidazole showing ∼60-80% efficacy in the acute stage. Unfortunately, benznidazole efficacy is extremely poor (∼7-20%) in the chronic stage (*6–8*). Similarly, nifurtimox treatment of CD patients resulted in decreased *T. cruzi* antibody levels and seroconversion in only a third of patients (*9*). There are no vaccines for CD, and its global burden well beyond endemic regions underscores the urgent need for chronic-stage treatments, such as therapeutic vaccines or new drugs.

During infection, non-replicative metacyclic trypomastigotes are released by the insect vector and enter the mammalian host, where they infect various cell types. Inside the cells, *T. cruzi* differentiates into replicative amastigotes. After several rounds of division, amastigotes differentiate into non-dividing trypomastigotes that disrupt host cells and are released into the bloodstream to infect new cells, unless they are taken up by the insect. The acute infection is marked by high blood parasitemia, which is typically controlled within the first weeks after infection. This is followed by low blood parasitemia and infection of cardiac, smooth, and skeletal muscles, adipose and other tissues; this multiple-tissue infection is a hallmark of chronic CD. Long-term tissue infection is often associated with inflammation of the heart and gastrointestinal tissues in symptomatic patients, leading to chronic disease. Antibodies and CD4^+^ and CD8^+^ T cells reduce parasite burden, but do not clear the infection (*10–13*). Parasites persist through mechanisms that are not yet fully understood. It has been postulated that changes in surface antigens expressed contribute to the parasites’ persistence by enabling them to evade antibodies and CD8^+^ T cells (*14*). About 0.2-7% of parasites seem to undergo dormancy in animal models, which could contribute to persistence upon drug treatment (*15*). Infection of immune-privileged tissues might also contribute to persistence due to restricted immune responses at those sites (*16–19*).

A bottleneck in developing a vaccine for CD is the lack of knowledge of vaccine targets, i.e., antigens that can stimulate antibody and cellular responses to control infection or disease burden without causing severe side effects or exacerbating the disease. A handful of antigens, typically selected by bioinformatic and experimental approaches (*20*) and delivered via adenoviruses (*21*), recombinant proteins (*22*), or mRNA (*23*) have been used, indicating the feasibility of vaccine development. This includes a glycotope that mimics surface protein glycosylation and induces α-Gal antibodies, immunizations with which reduced parasite burden in prophylactic vaccination of mice (*24*). Most tested antigens are members of large and diverse multigene families, such as trans-sialidases, mucins, and amastins (*20–22, 24*). Immunizations with these highly immunogenic antigens also reduced infection by inducing antibody production and CD8^+^ T cell expansion. However, *T. cruzi* strains are organized into several discrete typing units (DTUs) that exhibit significant diversity. There is extensive variability in the coding sequences of multigene family proteins within a strain and between DTUs, suggesting that these proteins are unlikely to be ideal vaccine targets. This is further underscored by recent findings on their expression variation each time parasites egress from a host cell (*14*). The presence of three distinct infectious forms, i.e., metacyclic, amastigotes, and trypomastigotes, poses additional constraints on vaccine target selection, since restricting antigen expression to a single infectious form could limit immune surveillance.

In this work, we sought to systematically identify *T. cruzi* proteins that are immunogenic in humans as potential vaccines and diagnostic candidates for CD. We generated a *T. cruzi* genome-wide library for yeast surface display (YSD) and screened for antigen-antibody binding using antibodies from patients with CD or healthy donors. We identified hundreds of *T. cruzi* antigens that elicit antibody responses in humans and defined their antibody-binding sequences. We also identified *T. cruzi* antigens conserved in humans, and their cross-reactivity with antibodies from healthy individuals suggests potential routes for the origin of autoantibodies.

Hundreds of antigens that induced *T. cruzi*-specific antibodies were conserved across DTUs, and quantitative proteomics revealed that several of them were expressed in all infectious forms of *T. cruzi*. Vaccination of infected mice with a yeast-based vaccine expressing a selected immunogenic sequence controlled *T. cruzi* infection from blood and tissues in both prophylactic and therapeutic models, demonstrating the approach’s efficacy and validating the antigen as a vaccine candidate. The results show a large dataset of antigens for vaccine, biomarker, and diagnostic development. It also shows that yeasts are a potential vaccine delivery system for CD and identifies a therapeutic vaccine candidate for chronic CD.

## RESULTS

### A *T. cruzi* genome-wide library for yeast surface display

We generated a genome-wide library of the *T. cruzi* Sylvio X10 strain for YSD (*25, 26*). In this system, *T. cruzi* proteins or protein fragments are expressed on the yeast surface via the Aga1p-Aga2p display system. *T. cruzi* DNA sequences were cloned in frame with Aga2p using the pYD1 vector, which contains a C-terminal Xpress epitope for tracking the fused proteins expressed on the yeast surface (**Fig. 1A**). The *T. cruzi*-Aga2p fused proteins are attached to the yeast surface via disulfide bonds to the membrane-associated Aga1p protein. As we reported previously, the Oxford Nanopore sequencing of the *T. cruzi* library revealed DNA fragments ranging from ∼0.3 to 3 kb, with a mean size of 1.1 kb (*25*). The library included 4.0×10^6^ clones, yielding ∼30-fold genome coverage, an average of 270 clones per gene, and included all coding sequences (*25*). We analyzed the library-predicted polypeptides from ∼7.2 million Nanopore sequences and found that they encoded 248,946 unique predicted polypeptides, yielding an estimated mean of 23.6 in-frame unique polypeptides per gene (*25, 26*). We transformed yeast and, to ensure whole-genome representation of *T. cruzi*, obtained clones sufficient to achieve ∼10-fold library coverage, yielding 5.0×10^7^ calculated yeast clones. Oxford nanopore sequencing of the yeast-transformed library revealed complete genome coverage (**Fig. 1B**). Comparison of the yeast-transformed library to the original library generated in bacteria showed a Pearson correlation coefficient of 0.91 (**Fig. 1C**), indicating that library representation was preserved after transformation and passages of the yeast. Western blot analysis using the Xpress epitope confirmed the galactose-induced expression of the library, with fragments migrating as multiple bands between 25 and 40 kDa, which is well within the expected range for most library protein length (**Fig. 1D**; Aga2p-Xpress has a calculated mass of 22 kDa, and its migration varies due to Aga2p glycosylation). Western analysis using antibodies against trypanosome phosphoglycerate kinase and peroxin 14 also confirmed *T. cruzi* protein expression in galactose-induced yeast lysate (**Fig. S1**), and yeast flow cytometry with CD patients’ serum antibodies showed over 70% antibody binding, confirming parasite protein expression (**Fig. S1**). Flow cytometry and immunofluorescence analysis also confirmed the surface expression of the proteins in >80% of the yeast (**Fig. 1E-F**). The data indicate a comprehensive *T. cruzi* YSD library and its functional expression in yeast.

**Fig. 1.**
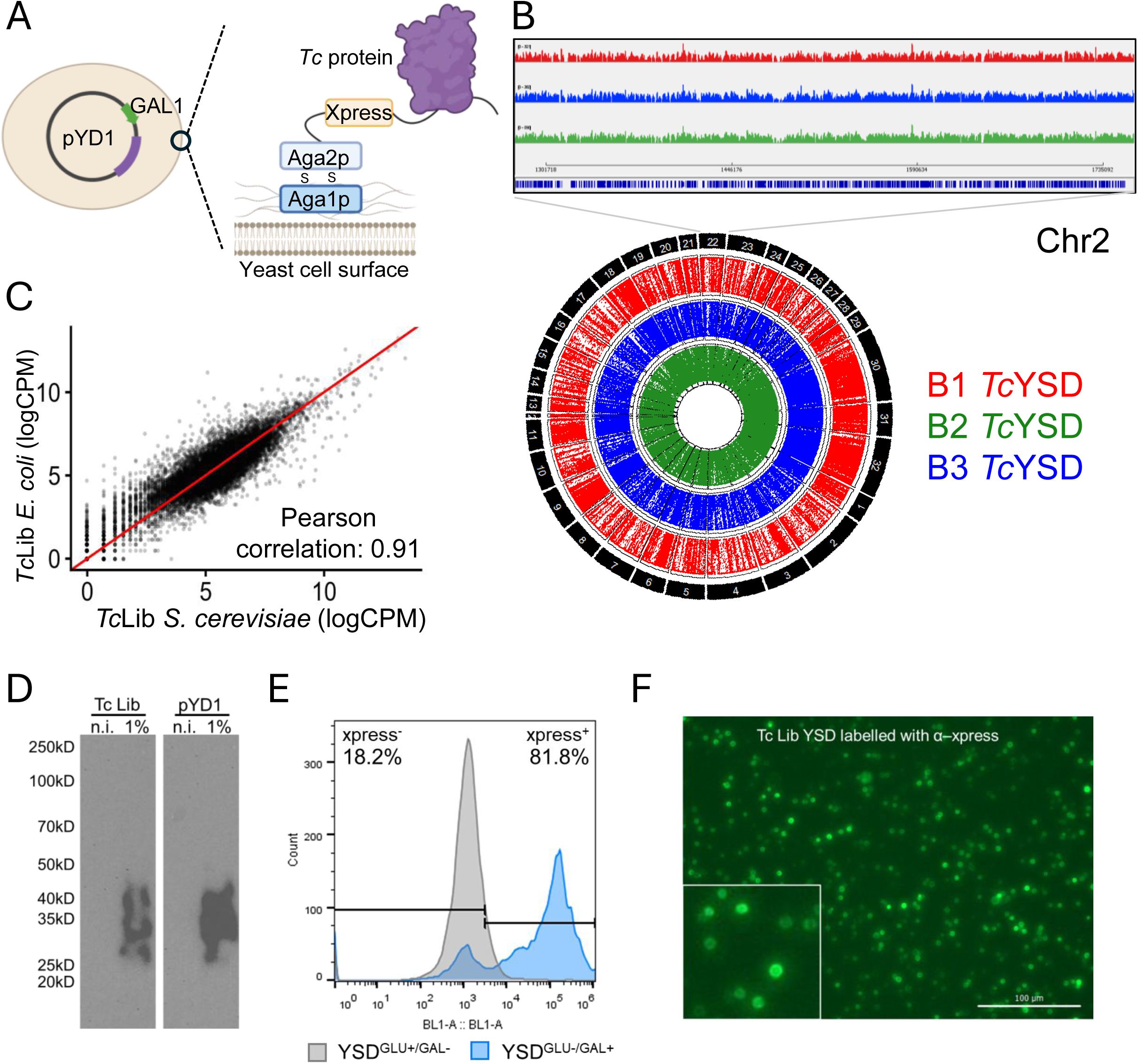
A *T. cruzi* genome-wide library for YSD. **A)** Diagram of YSD expressing a *T. cruzi* protein. The *T. cruzi* gene (or gene fragments) is cloned in frame with the yeast Aga2p protein. Aga2p interacts with Aga1p via disulphide bonds, resulting in the surface expression of the fused *Tc-*Aga2p protein. The *T. cruzi* proteins have a C-terminal Xpress epitope used to monitor protein expression. Created in BioRender. Cestari, I. (2026) https://BioRender.com/k3mul3q. **B)** Circular plot shows the mapping of *T. cruzi* genome-wide library extracted from yeast. Library DNA was sequenced by Oxford nanopore sequencing. B1, B2, and B3, biological replicates 1, 2 and 3. The inset indicates a segment of chromosome (Chr) 2, highlighting genome coverage. **C)** Correlation analysis of the *T. cruzi* genome-wide library after its generation in *Escherichia coli* (bacteria) and after transformation in *Saccharomyces cerevisiae* (yeast). **D)** Western blot of *S. cerevisiae* expressing Aga2p-Xpress only (empty pYD1 vector) or *T. cruzi* library induced to express the library with 1% galactose or to repress with 1% glucose (n.i., non-induced). The migration pattern of the pYD1 yeast reflects Aga2p glycosylation (*25*). **E-F)** Flow cytometry (E) and immunofluorescence (F) of yeast expressing *T. cruzi* library detected with α-Xpress antibodies. Bar = 100 µm.

### Identifying *T. cruzi* immunogenic antigens with Chagas disease patients’ antibodies

To identify *T. cruzi* proteins immunogenic in humans, we performed a screen using the *T. cruzi* YSD and pooled sera from patients with chronic CD or healthy individuals (HI). In this assay, the sera were incubated with the *T. cruzi* YSD library, and sera antibodies that bind *T. cruzi* antigens expressed on the yeast surface were used to select yeast binders via magnetic-activated cell sorting (MACS) with Protein-G magnetic beads (**Fig. 2A**). The bound yeast was eluted, re-grown, and the procedure was repeated a total of three times to enrich the yeast population. The library DNAs were extracted from yeast and sequenced using Oxford nanopore sequencing.

**Fig. 2.**
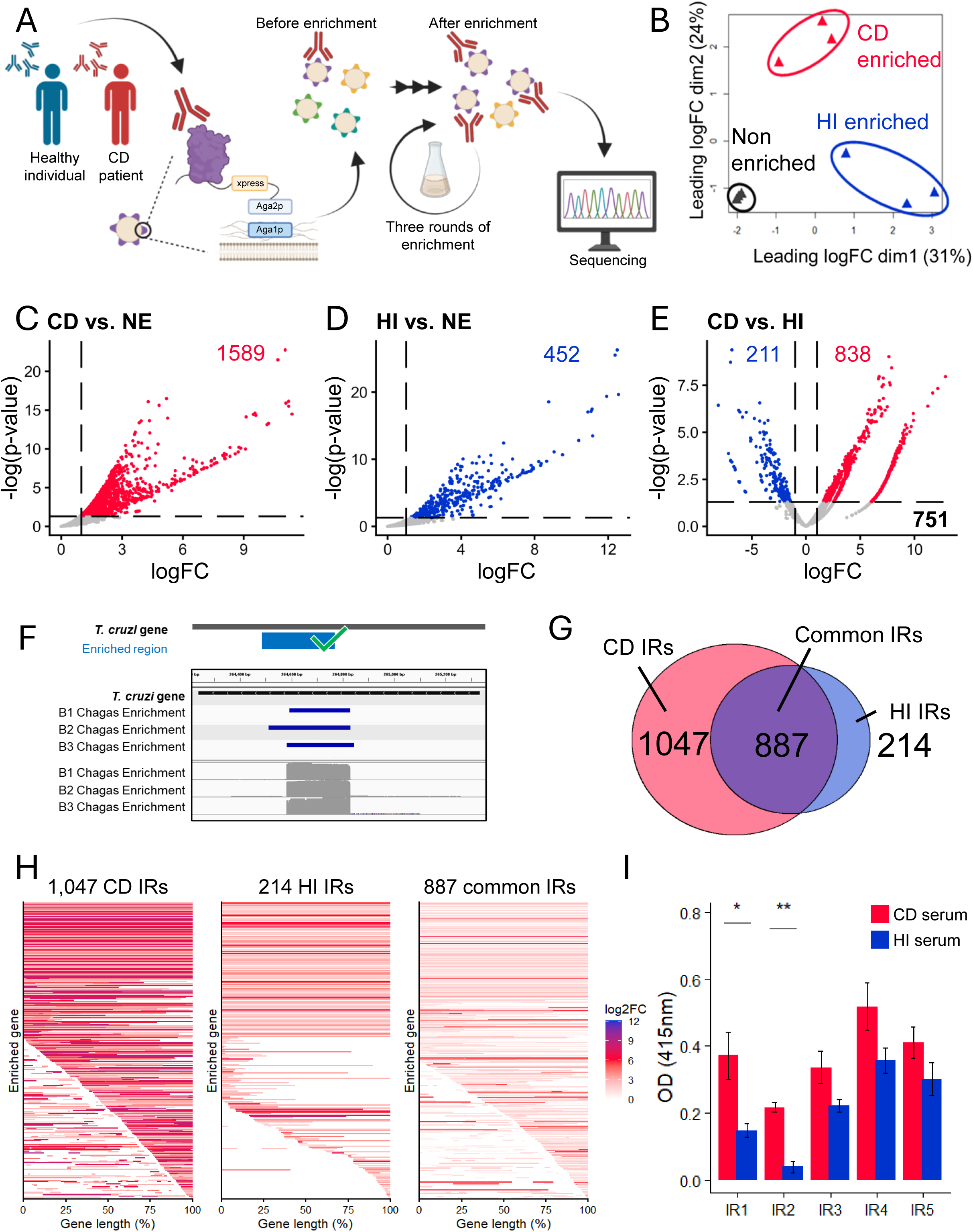
YSD screen with patient antibodies identifies *T. cruzi* immunogenic sequences. **A)** Diagram of the *T. cruzi* YSD library screen with CD patients’ antibodies. Antibodies from CD patients or HI were mixed with yeast expressing *T. cruzi* proteins. After three rounds of MACS enrichment, library DNA was extracted from yeast for Oxford Nanopore sequencing. Created in BioRender. Cestari, I. (2026) https://BioRender.com/7b0mssr. **B)** PCA of *T. cruzi* YSD libraries enriched with CD patients’ antibodies, HI antibodies, or non-enriched. **C-E)** Volcano plots of *T. cruzi* YSD library enriched with (C) CD pooled sera compared to the non-enriched library, (D) HI’s pooled sera compared to the non-enriched library, and (E) the comparison of statistically significant (fold-change ≥ 2, *p-*value ≤ 0.05) CD-enriched genes (from C) to HI-enriched genes (from D). **F)** Diagram shows Oxford nanopore reads enrichment of *T. cruzi* YSD library, indicating DNA sequence encoding immunogenic region (IR), i.e., antibody-binding site. B1-B3, biological replicates 1-3. **G)** Venn diagram of the number of IRs identified exclusively with CD or HI sera antibodies, or common to both. **H)** IRs of *T. cruzi* YSD library exclusively enriched with antibodies from CD patients or HI, or IRs common to both. The coloured regions indicate significantly enriched IR (log2 fold-change, FC). **I)** ELISA with CD patients or HI sera binding to yeast surface expressing *T. cruzi* selected IRs. IR1 (TcSx10.chr5.027000); IR2 (TcSx10.chr8.042270), IR3 (TcSx10.chr5.027000, different region with lower enrichment), IR4 (TcSx10.chr5.028600), IR5 (TcSx10.chr27.112540).

Principal component analysis (PCA) revealed differences between non-enriched and enriched libraries for antibodies from CD patients and HI donors, and consistency across biological replicates (**Fig. 2B**). We identified 1,589 enriched *T. cruzi* proteins reacting with CD patients’ antibodies (fold-change ≥ 2-fold, p-value ≤ 0.05) compared to the library before antibody binding selection (**Fig. 2C, Data file S1**). There were 452 *T. cruzi*-enriched (fold-change ≥ 2-fold, *p*-value ≤ 0.05) proteins with HI donors’ antibodies (**Fig. 2D, Data file S1**), revealing the presence of antibodies cross-reacting with *T. cruzi* proteins. Comparison of the statistically significant hits from CD and HI antibody screens revealed 838 *T. cruzi* antigens (fold-change ≥ 2-fold, p-value ≤ 0.05) that specifically reacted with CD patient antibodies (**Fig. 2E, Data file S1**), representing a pool of *T. cruzi* proteins that are immunogenic in humans and elicit *T. cruzi-*specific antibody responses. 751 *T. cruzi* antigens reacted with both CD and HI sera antibodies, and 211 *T. cruzi* antigens reacted only with antibodies from HI, suggesting the presence of cross-reacting antibodies against the *T. cruzi* antigens (**Fig. 2E**), perhaps due to unrelated infections or vaccinations. Because the library was designed with a range of DNA sequence lengths, the antibody reaction reveals specific antibody-binding sites at nucleotide resolution within *T. cruzi* antigens (**Fig. 2F**), indicating one or more immunogenic regions (IRs) within the identified proteins (**Fig. 2G-H**). Notably, enrichment with CD-specific antibodies was greater than with HI or the common antigen subset (**Fig. 2H**). To validate the binding reactions, we generated yeast cells expressing the IRs of five antigens selected using CD-specific antibodies and performed ELISAs with sera from CD patients or HI. The data showed greater enrichment with CD antibodies than with HI antibodies (**Fig. 2I**), confirming their immunogenicity and ruling out potential screen artifacts. The reactivity of HI sera in the ELISA reflects unspecific antibodies cross-reacting with the yeast. The screen identified hundreds of *T. cruzi* antigens that elicited antibody responses in humans and their IRs. The identification of parasite-specific antigens and those conserved with host proteins may help identify vaccine targets and antigens inducing autoantibodies, respectively. The data provide insights into the *T. cruzi*-specific antibody response in patients with chronic CD.

### Immunogenic antigens expression and reactivity across CD patients

Because *T. cruzi* has three infectious stages that circulate in the host, we sought to determine which stages express the antigens identified in the YSD screen by investigating their stage-specific expression. We generated tandem mass tag labelling and mass spectrometry (TMT-MS) of whole-cell metacyclic trypomastigotes (MT), column-purified amastigotes (AM) from infected cardiomyocytes (H9c2 cells), and cell (H9c2)-derived trypomastigotes (CT), quantitatively covering 3,194 stage-specific *T. cruzi* proteins (*14*). The data showed that 447 IR-containing proteins detected by TMT-MS were expressed among all three *T. cruzi* infectious stages (**Data file S2**). Of those, 235 IRs were CD-specific, 173 were common between CD and HI, and 39 were specific to HI with over 95% of them expressed across the three infectious stages (**Fig. 3A, Fig. S2**). Because the TMT-MS did not cover all *T. cruzi* proteins, other IRs were benchmarked using RNA-seq data, confirming their expression in CTs, AMs, or MTs (**Fig 3A**, **Fig. S2**). Notably, of the 838 CD-specific antigens (**Fig. 2E**), only ∼20% represented genes encoding members of multigene families, such as trans-sialidases and mucins, whereas ∼42% lacked functional annotation (hypothetical proteins), and the remaining ∼38% included annotated proteins (**Data file S1**). Similarly, common proteins between CD and HI included ∼22% multigene family genes, ∼38% hypothetical proteins, and ∼40% with functional annotation. The HI-specific antigens contained 28% of multigene family genes, which may reflect cross-reacting antibodies against common carbohydrate domains within this protein family, as well as proteins that are typically highly conserved across organisms, such as histones, helicases, kinases, and ABC transporters (**Data file S1**).

**Fig. 3.**
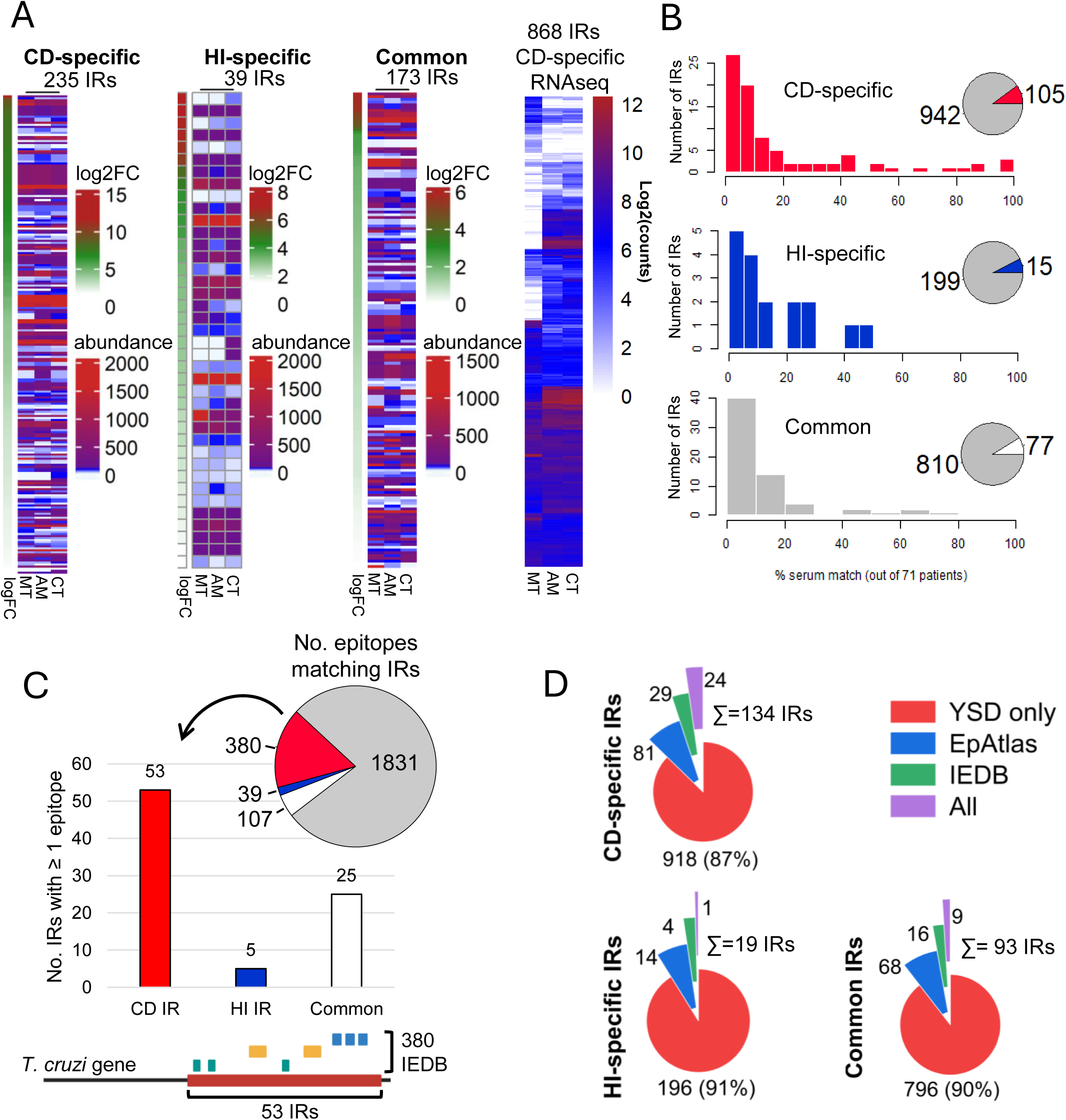
Expression of IR-containing proteins and immunogenicity across CD patients. **A)** Expression of IR-containing proteins in *T. cruzi* metacyclic trypomastigotes (MT), amastigotes (AM), and cell-derived trypomastigotes (CT) detected by TMT-MS analysis. IRs identified in CD or HI screens, or those common to both, are shown. The data show the log2 fold change (FC) in YSD enrichment and protein abundance as determined by TMT-MS. Of CD-enriched IRs, RNA-seq of gene expression is shown (right). Abundance (log2) of normalized counts per million is shown. **B)** Distribution of IR sequences overlapping with peptide sequences matching those in the EpAtlas. Inset pie plots show the number of IRs that match peptides in each group (red, blue, white) and those that do not (grey). **C)** Number of IRs with sequences matching epitopes from IEDB. The inset pie plot shows the number of epitopes that match IR sequences out of the total epitopes analyzed. The IR can contain one or more epitopes, as indicated in the diagram below. **D)** Number of IRs that react with serum from CD patients, HI, or both (Common), and the aggregated number of epitopes from the EpAtlas and IEDB. All, IR regions matching both IEDB and the EpAtlas.

To further validate the identified antigens, we compared our dataset with 3,868 *T. cruzi* peptides screened against sera from 71 CD patients (EpAtlas) (*27*). We identified 152 unique peptides matching 197 IRs enriched in the YSD screen (**Fig. 3B**). Of the 197 IRs, 105 included IRs that reacted specifically with antibodies from CD patients, 77 were shared between CD and HI, and 15 IRs were those identified specifically with HI sera (**Fig. 3B, Data file S3**). Notably, 15.2% of these IR sequences matched peptides recognized by antibodies from at least 30 patients, indicating that several sequences were immunogenic across the patients. Furthermore, we analyzed our dataset against the Immune Epitope Database (IEDB) (*28*) for additional *T. cruzi* antigens known to react with patients’ antibodies. We found 526 IEDB unique peptides matching 83 unique IRs, of which 53, 25, and 5 were CD-specific, cross-reacted with CD and HI sera, and HI-specific, respectively (**Fig 3C**). Overall, the analysis identified 134 IRs that matched peptides reacting with individual sera from CD patients (**Fig. 3D**), whereas 93 IR-containing matching peptides cross-reacted with both CD and HI sera. The sequence overlap between peptides and IRs supports the immunogenicity of IRs across patients. Lastly, 19 IR-containing peptides reacted with HI sera, likely reflecting false positives in the peptide dataset. Our dataset identified 918 additional CD-specific IRs not reported in previous studies, reflecting antibodies that recognize folded antigens (rather than linear peptides) and post-translational modifications. The data identified antigens that react with antibodies prevalent in the chronic stage of CD and those that cross-react with HI, revealing the presence of conformational immunogens with potential therapeutic and diagnostic applications.

### Immunogenic regions as vaccine targets for Chagas disease

We postulated that some IRs are potential vaccine targets against *T. cruzi*. To identify and prioritize IRs with vaccine target profiles, we searched for (i) top enriched IRs (FC ≥ 2, *p*-value ≤ 0.05) reacting exclusively with CD patients’ antibodies (i.e., 1,047 IRs, **Fig. 2G**), (ii) IRs that are conserved across *T. cruzi* DTUs, (iii) IRs that are not part of the variable large multigene families, (iv) nor conserved in humans/mice, and (v) IRs with evidence of protein expression across all three *T. cruzi* infectious stages. We analyzed the 1,047 CD-specific IRs across several *T. cruzi* genomes and identified 917 IRs that were conserved across DTUs I, II, III, IV, V, and VI, all of which include strains infectious to humans (*29*) (**Fig. 4A-B**). We excluded 157 proteins conserved in humans, resulting in 760 non-human homologs (**Fig. 4B, Data File S4**). This dataset contained 263 IRs from variant multigene family proteins (e.g., trans-sialidases, mucins, MASPs), and after their removal, we retained 497 conserved IRs. Of these, 143 IRs were from proteins detected by TMT-MS to be expressed across all *T. cruzi* infectious forms, i.e. MTs, AMs, and CTs (**Fig. 4B**). The immunogenicity, conservation across DTUs, and TMT-MS evidence of protein expression indicate that at least 143 IRs are potential candidates for vaccine or CD-specific diagnostic studies, and this number could increase with further proteomic evidence of their expression across the three parasite infectious forms. Gene ontology (GO) enrichment analysis of this protein subset indicated that most proteins lacked functional annotation (i.e., hypothetical protein), whereas the remaining were membrane proteins, intracellular proteins, or enzymes (**Fig. 4C)**.

**Fig. 4.**
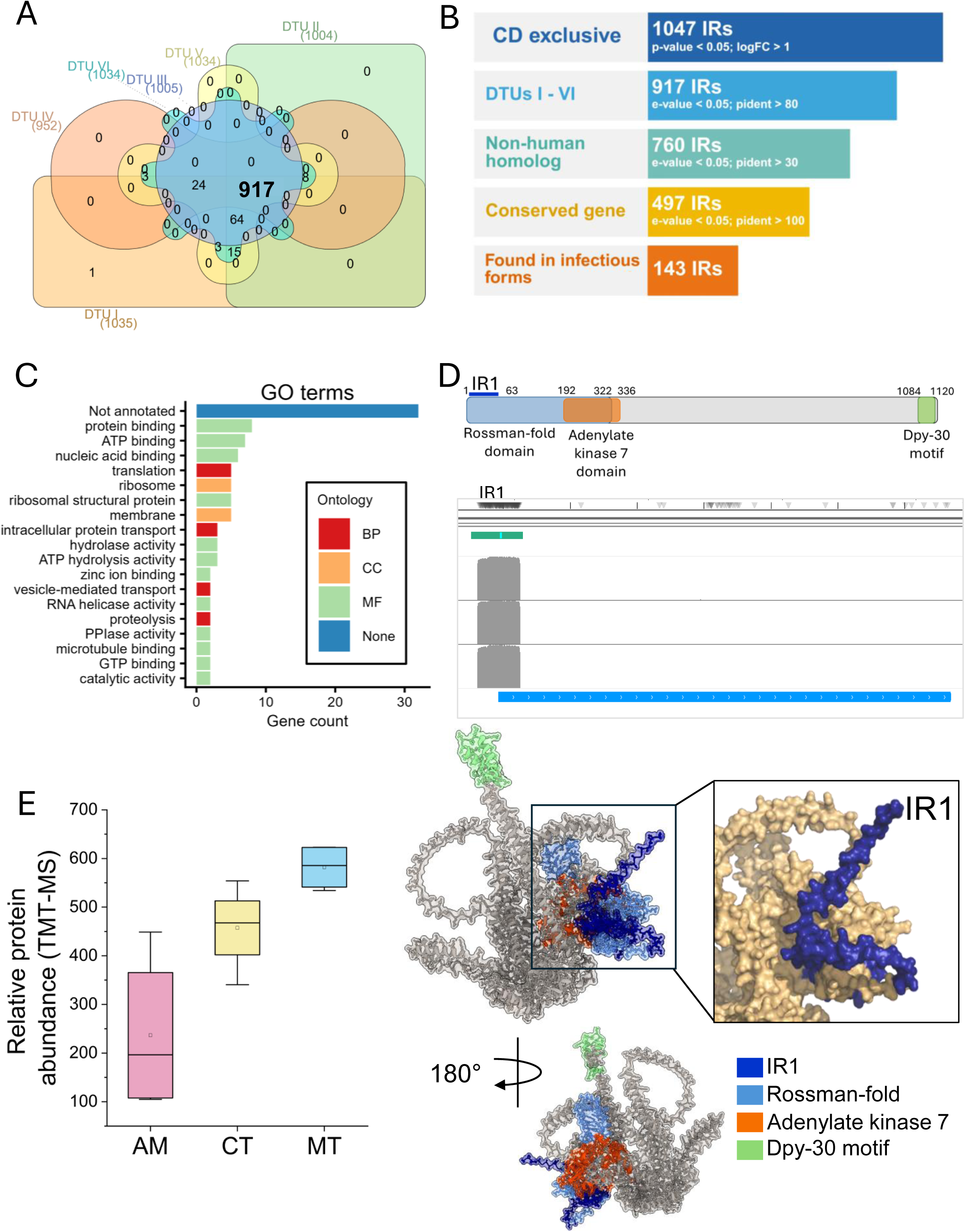
Prioritizing vaccine candidates for Chagas disease. **A)** The Venn diagram shows the number of CD-specific IR sequences conserved or unique to *T. cruzi* discrete typing units (DTUs). **B)** The priority diagram shows the changes in the number of CD-specific IRs after the removal of i) non-conserved sequences among DTUs, ii) human homologs, iii) multigene family members, iv) those without evidence of expression in all three infectious stages (MT, AM, CT). Created in BioRender. Cestari, I. (2026) https://BioRender.com/g23vlci. **C)** Gene ontology (GO) enrichment analysis of the selected 143 IRs (from B). BP, biological processes; CC, cellular components; MF, molecular functions. **D)** In the top, a diagram of the protein-encoding selected IR1 showing its domains. Middle, Oxford Nanopore reads enrichment of IR1 mapping on the coding sequence (GenBank ID: TcSx10.chr5.027000). Below, AlphaFold 3 structure prediction of the hypothetical protein encoding IR1. The IR1 region corresponding to 63 amino acids at the protein N-terminus is shown in blue. **E)** Abundance IR1-encoding protein as detected by TMT-MS in the *T. cruzi* infectious forms. AM, amastigotes; CT, cell-derived trypomastigotes; MT, metacyclic trypomastigotes.

We selected one of the top-ranked IRs, corresponding to a 63 aa at the N-terminus of a hypothetical protein (Gene ID: TcSx10.chr5.027000 from *T. cruzi* Sylvio X10 strain genome, GenBank access # JBMETK000000000.1). The encoded protein in *T. cruzi,* as well as its ortholog in *T. brucei,* was found in flagellar fractions by mass spectrometry analysis (*30, 31*), suggesting that it is likely a flagellar protein. Enrichment analysis of the three biological replicates consistently identified the IR1 at the N-terminus of the protein, as demonstrated by the piled-up reads predominating at 5’-end of the coding sequence (**Fig. 4D).** The protein has an N-terminal NAD(P)-binding Rossmann-fold domain (aa 1-322), an adenylate kinase 7 domain (aa 192-336), and a C-terminal Dpy-30 motif (aa 1084-1120), with the N-terminal IR1 predicted to be surface-exposed in the AlphaFold 3 structure (**Fig. 4D**, see **Fig. S3** for protein sequence and AlphaFold 3 metrics). Quantitative proteomics by TMT-MS showed expression of the IR1-encoding protein in MT, AM, and CT (**Fig. 4E**), with high levels in trypomastigote forms (MT and CT). The expression of the protein encoding the IR1 sequence in the *T. cruzi* infectious forms, its conservation across DTUs, and its immunogenicity suggest it is a potential vaccine target.

### Immunizations with yeast-delivered IR1 control *T. cruzi* infection in mice

Since *T. cruzi* naturally infects rodents, we evaluated whether prophylactic IR1 vaccinations could reduce infection burden in a murine CD model (**Fig. 5A**). In mice, the acute stage lasts ∼40 days and is characterized by a high parasite load, followed by a chronic stage lasting several months, with subpatent parasitemia and persistent tissue infections (11, 12). First, we expressed 116-aa N-terminal sequence of the protein-coding IR1 (from M1 to D116) and purified it as a His-tagged recombinant IR1 (rIR1) from *E. coli* (**Fig. 5B**). rIR1 migrated predominantly at ∼12.7 kDa, with minor protein amounts migrating as dimers/oligomers, as confirmed by α-His Western blot and proteomic analysis (**Fig. 5B, Fig. S4**). We reasoned that inactivated baker’s yeast (*S. cerevisiae*) might be an efficient vehicle for vaccination, as it is non-infectious and can express high levels of the target antigen on its surface. We generated yeast expressing IR1 (yIR1) or the empty vector pYD1 (y, as a control) and confirmed protein expression in yeast using α-Xpress antibodies (**Fig. 5B**). Antibody-binding assays also confirmed yIR1 reactivity with patients’ sera antibodies (**Fig. 2I**). Yeast expressing IR1 was heat-inactivated (60°C for 90 min), lyophilized, and stored at -80°C. Immediately before immunizations, yIR1 was hydrated in PBS, and its protein integrity and surface expression were assessed by Western blot and flow cytometry, revealing surface expression in >70% of yeast cells (**Fig. 5B-C, Fig. S5**).

**Fig. 5.**
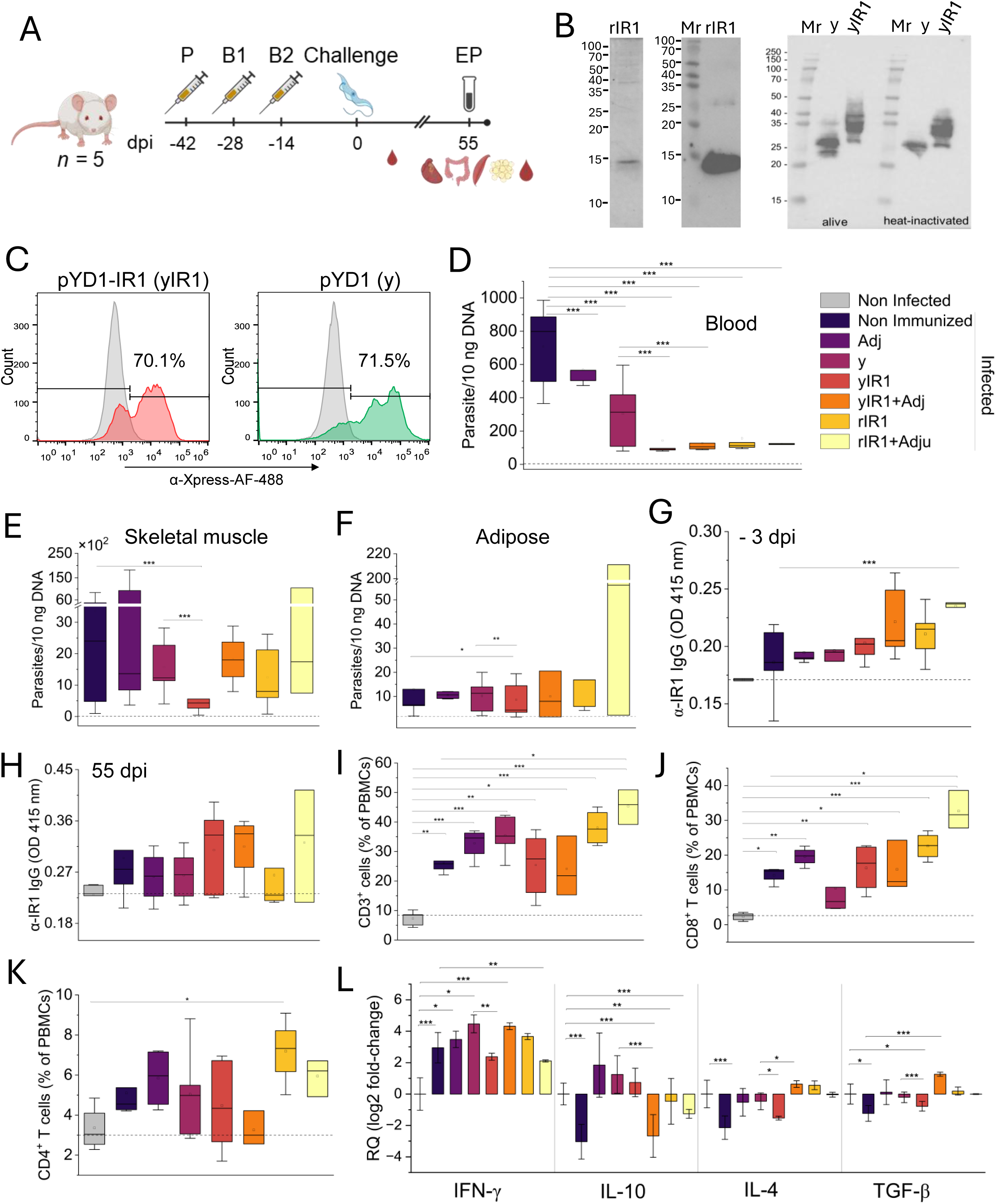
Prophylactic vaccination of mice with rIR1 and yeast-delivered IR1. **A)** Diagram of mice prophylactic vaccination. P, prime; B, boost; EP, endpoint. Blood was collected at multiple time points, and organs were collected at the endpoint. Day 0 corresponds to the day mice were infected. Created in BioRender. Cestari, I. (2026) https://BioRender.com/8lje6kw. **B)** Left, Coomassie-stained 15% SDS/PAGE of His-tagged IR1 purified from *E. coli* (rIR1). Middle, Western blot with α-His antibodies of rIR1 resolved in 15% SDS/PAGE. Right, Western blot with α-Xpress antibodies of yeast expressing pYD1 (empty vector, y) or pYDI-IR1 (yIR1). Lysates were prepared from fresh yeast (alive) or from lyophilized yeast (heat-inactivated at 60 °C for 90 min, then lyophilized, stored at -80 °C, and rehydrated in PBS). The pYD1 vector expresses the 22 kDa Aga2p-Xpress epitope. **C)** Flow cytometry analysis of yeast expressing pYD1(y) or pYD1-IR1 (yIR1) with anti-Xpress antibodies after heat-inactivation at 60 °C for 90 min. **D)** Quantification of blood parasitemia in vaccinated and/or infected mice by qPCR of DNA at 28 dpi (parasitemia peak). **E-F)** Quantification of parasite load in hindlimb skeletal muscle tissue (E) or adipose tissue (F) by qPCR of DNA. The dotted horizontal line indicates the qPCR noise levels established from non-infected mice for each tissue. **G-H)** Quantification of α-IR1 antibodies by ELISA from mice 3 days before (G) and at the end point (55 dpi) of infection (H). ELISA plates were coated with yeast isolated IR1. **I)** Percentage of T cells in blood detected by flow cytometry using α-CD3 antibodies. **J-K)** Frequency of CD8^+^ T cells (CD3^+^CD8^+^) (J) or CD4^+^ T cells (CD3^+^CD4^+^) (K) from total peripheral blood mononuclear cells (PBMCs) from mice detected by flow cytometry. **L)** Expression of cytokines from mice PBMCs by qPCR of cDNAs. Data show mean ± standard error of the mean (SEM). For D to L, n = 5 mice per condition. For box plots, the horizontal line within the box indicates the median, the box represents the interquartile range (IQR, 25th–75th percentiles), and the whiskers extend to 1.5 times the IQR. The open square within the plot shows the mean. Individual points represent outliers. For graphs in H to L, the dotted line shows the median values of non-infected mice. *, *p* ≤ 0.05, ** *p* ≤ 0.01, *** *p* ≤ 0.001. Nonparametric multiple-contrast test with Dunnett-type contrasts was used to compare experimental groups against the non-immunized group.

We performed three rounds of immunizations in mice, with two boosts at 7-day intervals after 14 days of priming (**Fig. 5A**). We used IL-12 and GM-CSF as adjuvants, which stimulate Th1-type T-cell responses (*32–34*), including in CD vaccine experiments (*35, 36*). Mice were challenged 14 days after the last immunization with 10^5^ MTs of *T. cruzi* Sylvio X10 strain.

Parasite load in blood and tissues was analyzed by real-time quantitative PCR (qPCR) throughout the infection and at the endpoint of the experiment (55 days post-infection, dpi). We confirmed that all mice were infected (**Table S1**), and the infection or vaccinations did not significantly affect mouse weight (**Fig. S6**). There was a significant decrease in *T. cruzi* load from blood in mice vaccinated with yIR1 or rIR1 in the presence or absence of adjuvants at the parasitemia peak (28 dpi) (**Fig. 5D**, *dotted baseline indicates mean qPCR values of non-infected mice used to define the background noise*). Low levels of parasites were also detected in the blood at the endpoint (55 dpi) (**Table S2**). Because blood parasitemia is typically low at this point (55 dpi) due to the mice’s natural immune response, the low parasite levels may reflect DNA from dead parasites or parasites emerging from tissues. Vaccinations with yeast expressing the empty vector pYD1 (y) also caused a decrease in *T. cruzi* infection, suggesting that yeast alone may have an adjuvant effect (**Fig. 5D**). Notably, there was a significant decrease in *T. cruzi* load in the muscle tissues of yIR1-immunized mice approaching that of non-infected mice (**Fig. 5E**). In contrast to what was observed in blood, rIR1 with or without adjuvants did not provide tissue protection, whereas the rIR1/Adj seemed to have a detrimental effect on tissue infection (**Fig. 5E-F**). Because not all mice had detectable organ infections or exhibited a variable or low-level infection (**Table S2**), comparative analysis of other tissues was inconclusive, although a significant two-fold decrease was observed with yIR1 in adipose tissue (**Fig. 5F**). The data suggest that yeast-delivered IR1 is a more effective vaccine than its recombinant form, possibly due to a combination of high IR1 surface expression in yeast, protein folding, potential IR1 post-translational modifications, and a potential yeast adjuvant effect (*37*).

Antibody analysis showed a trend toward increased α-IR1 antibody levels in mice vaccinated with yIR1, with or without adjuvants (**Fig. 5G-H**; *the dotted horizontal line indicates the median values in non-infected mice)*. In contrast, rIR1 vaccinations enhanced antibody production primarily in the presence of adjuvants. The T-cell population in the peripheral blood of both infected and vaccinated/challenged mice also increased (**Fig. 5I**). This increase reflects the expansion of cellular defenses, including CD8^+^ and CD4^+^ T cells (**Fig. 5J-K**), of which ∼95% were effector memory cells (CD8^+^CD44^hi^CD62L^lo^ and CD4^+^CD44^hi^CD62L^lo^) (**Fig. S7**), suggesting that yIR1 and rIR1 vaccinations can induce memory T cells. While these responses were effective in yIR1 to control parasite blood and tissue infection, the increased levels of α-IR1 antibodies and T cells in rRI1/Adj vaccinations did not correlate with tissue protection (**Fig. 5E-F**). There was increased expression of interferon-γ (IFN-γ) in PBMCs from infected and vaccinated mice (**Fig. 5L**), which was also reflected in IFN-γ serum levels (**Fig. S8**), although mice that received adjuvants had lower IFN-γ serum levels than yIR1 and rIR1 alone. INF-γ is required for activating parasite-killing responses (*38*). Their low levels in the presence of high CD8^+^ T cells in rIR1/Adj suggest an inefficient CD8^+^ T cell population, reflected in limited tissue protection (**Fig. 5E-F)**. This could be related to CD8^+^ T cell exhaustion, which has been reported in *T. cruzi* infection (*39*). To evaluate the anti-inflammatory response, we analyzed expression of interleukin-10 (IL-10), IL-4, and transforming growth factor-beta (TGF-β) in PBMCs (**Fig. 5L)**. IL-10, IL-4, and TGF-β transcripts were downregulated in non-vaccinated and infected mice, consistent with low anti-inflammatory responses during early stages of *T. cruzi* infection (*40*).

However, yIR1-vaccinated mice showed near-basal anti-inflammatory cytokine expression levels comparable to non-infected, non-vaccinated controls, suggesting a restored anti-inflammatory response to counterbalance the immune response to infection. This homeostatic control aligns with the effector response induced by yIR1 vaccination, which drove elevated IFN-γ levels, increased T cell counts, and a concomitant reduction in blood and tissue parasitemia (**Fig. 5D-F)**. The data suggest that yIR1 prophylactic vaccinations have protective and adjuvant effects controlling *T. cruzi* infection in blood and muscle tissues.

### Yeast-based IR1 therapeutic vaccination controls *T. cruzi* chronic infection

Because *T. cruzi* infections cause long-term chronic disease and have spread in non-endemic regions, we evaluated whether IR1 vaccinations could reduce infection burden in a therapeutic vaccine model in which mice are infected and then treated by vaccinations at the chronic stage (**Fig. 6A**). We infected mice with 10^5^ MTs of *T. cruzi* Sylvio X10 strain, which recapitulates chronic CD in mice (*41, 42*), and confirmed infection by qPCR of peripheral blood at 5 dpi, demonstrating that all mice were infected (**Table S3**). We treated infected mice with yIR1 or yeast alone (y, expressing the empty pYD1 vector) starting at 55 dpi. We performed three vaccinations at 7-day intervals, with or without IL-12 and GM-CSF as adjuvants and performed qPCR on blood and tissues at 100 dpi (endpoint). Infection or vaccinations did not significantly affect the mice’s weight (**Fig. S6**). We found complete *T. cruzi* clearance from muscle tissues in mice vaccinated with yIR1 (**Fig. 6B).** There was also a significant decrease (118-fold) of parasites from the intestine in yIR1-vaccinated mice approaching clearance (**Fig. 6C)**, whereas blood and other tissues retained low or variable levels of detection (**Table S4**), and with a two-fold decrease in adipose tissue (**Fig. 6D**). The low load may reflect persistent parasites or DNA amplification of dead parasites, suggesting that a longitudinal analysis may be required to evaluate clearance on these tissues. Notably, the addition of adjuvants did not increase protection compared to yIR1 alone (**Fig. 6B**), analogous to what was observed in the prophylactic model (**Fig. 5E-F**). The protective effect of yIR1 correlated with high α-IR1 antibody levels (**Fig. 6E**), suggesting that α-IR1 antibodies may have contributed to protection after vaccination. There was also an increase in peripheral CD8^+^ and CD4^+^ T cells in yIR1-immunized mice (**Fig. 6F-H**), of which ∼95% were effector memory T cells (CD8^+^CD44^hi^CD62L^lo^ and CD4^+^CD44^hi^CD62L^lo^) (**Fig. S7**). Gene expression analysis in PBMCs showed increased IFN-γ levels after infection or vaccination (**Fig. 6I**) and slightly higher serum levels in yIR1-vaccinated mice (**Fig. S8**). In contrast to the acute stage, the anti-inflammatory response increased in non-vaccinated mice (**Fig. 6I**), potentially counterbalancing the immune response and allowing parasites to persist. But yIR1-vaccinated mice showed baseline-to-slightly low levels of IL-4 and TGF-β, suggesting a balanced immune response. IL-10 levels were below the limit of detection by ELISA or qPCR. As observed in the prophylactic model, the high level of CD8^+^ T cells in mice vaccinated with yIR1/Adj did not translate into greater protection, suggesting that this adjuvant (IL-12/GM-CSF) had a detrimental effect on the immune response, in contrast to the protective effect of yIR1 alone. The data show that inactivated yeast can serve as a vehicle for delivering a CD vaccine antigen. Moreover, it shows that yIR1 vaccination effectively clears muscle infection and reduces infection burden in mice, making it a potential therapeutic vaccine candidate for CD.

**Fig. 6.**
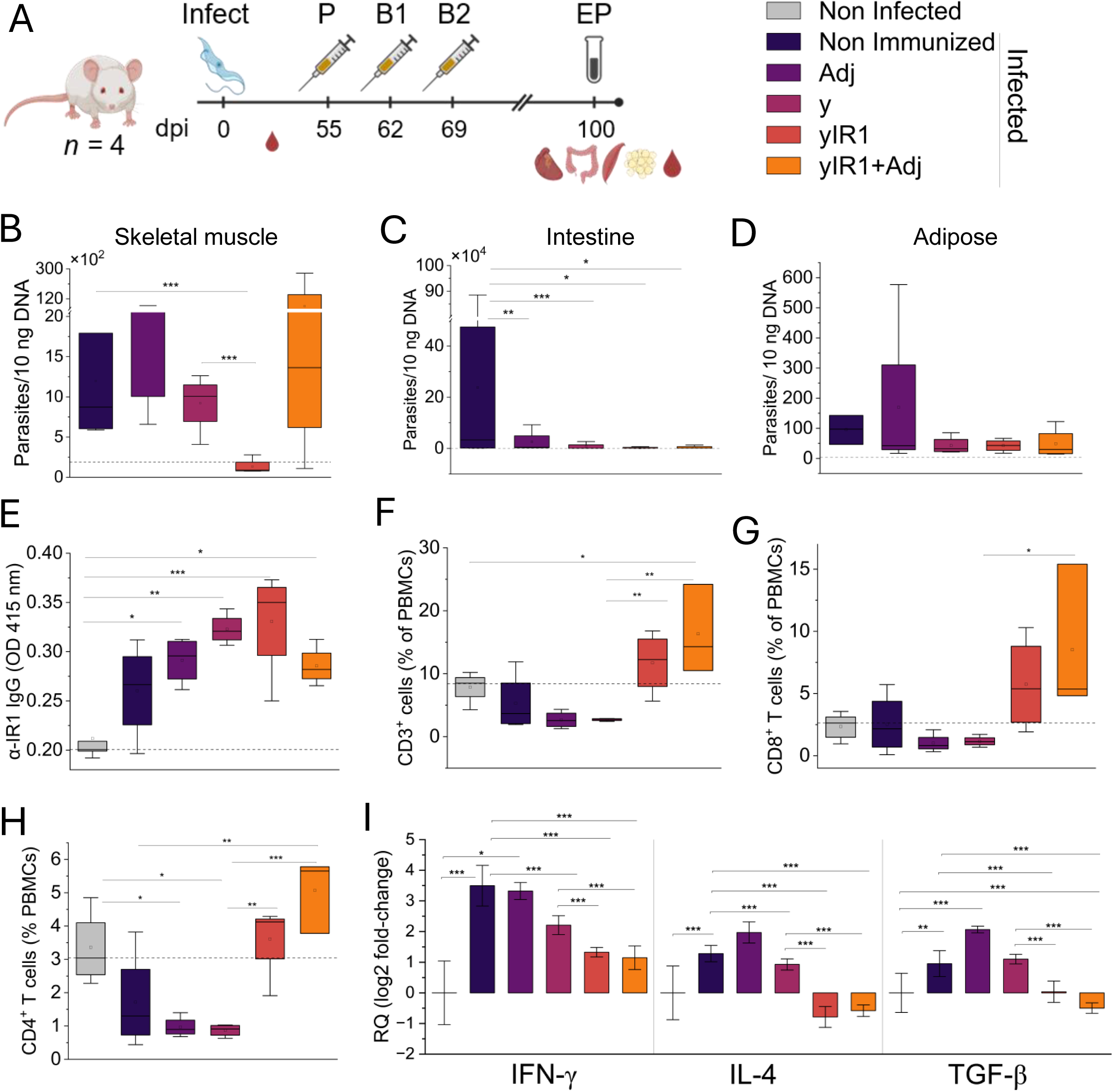
Therapeutic vaccination of mice with yeast-delivered IR1. **A)** Diagram of mice therapeutic vaccination. P, prime; B, boost; EP, endpoint. Blood was collected at multiple time points, and organs were collected at the endpoint. Day 0 corresponds to the day mice were infected. Created in BioRender. Cestari, I. (2026) https://BioRender.com/8lje6kw. **B-D)** Quantification of parasite load by qPCR of DNA in hindlimb skeletal muscle (B), intestine (C), and adipose tissues at 100 dpi (endpoint). The dotted horizontal line indicates the qPCR noise levels established from non-infected mice for each tissue. **E)** Quantification of α-IR1 antibodies by ELISA from mouse sera at 100 dpi. ELISA plates were coated with yeast isolated IR1. **F)** Percentage of T cells in blood detected by flow cytometry using α-CD3 antibodies. **G-H)** Frequency of CD8^+^ T cells (CD3^+^CD8^+^) (G) or CD4^+^ T cells (CD3^+^CD4^+^) (H) from mice’s PBMCs by flow cytometry. **I)** Expression of cytokines from mice PBMCs by qPCR of cDNAs. Data show mean ± SEM. For B to I, n = 4 mice per condition. For box plots, the horizontal line within the box indicates the median, the box represents the interquartile range (IQR, 25th–75th percentiles), and the whiskers extend to 1.5 times the IQR. The open square within the plot shows the mean. Individual points represent outliers. For graphs E-H, the dotted line shows the median values for non-infected mice. *, *p* ≤ 0.05, ** *p* ≤ 0.01, *** *p* ≤ 0.001. Nonparametric multiple-contrast test with Dunnett-type contrasts was used to compare experimental groups against the non-immunized group.

**Table 1.** Top 25 CD-specific IRs. Enrichment was calculated by comparing statistically significant CD-enriched IRs to HI-enriched IRs to define CD-specific IRs. Log2FC, log2 fold change; log2CPM, log2 counts per million. IR_ID, IR identification, which combines gene ID with the IR genome starting position. For a complete list and IR sequences, see Data File 4.

| IR_ID | log2<br>FC | log2CP<br>M | p-value | Annotation |
| --- | --- | --- | --- | --- |
| TcSx10.chr18.082030_31850 | 11.21 | 14.52 | 3.19E-16 | surface antigen 2 (CA-2) putative |
| TcSx10.chr2.004550_266650 | 8.32 | 13.54 | 6.62E-11 | calpain-like protein putative |
| TcSx10.chr5.027000_166962 | 7.05 | 11.44 | 2.79E-08 | hypothetical protein |
| TcSx10.chr30.127060_879550 | 8.05 | 10.93 | 1.68E-09 | monoglyceride lipase putative |
| TcSx10.chr4.022790_967600 | 7.63 | 10.74 | 6.34E-09 | hypothetical protein |
| TcSx10.chr27.112540_234200 | 7.54 | 10.71 | 8.52E-09 | flagellum-adhesion glycoprotein putative |
| TcSx10.chr2.007290_912050 | 5.95 | 10.42 | 1.04E-06 | hypothetical protein |
| TcSx10.chr2.007290_912250 | 5.95 | 10.42 | 1.04E-06 | hypothetical protein |
| TcSx10.chr2.007290_915150 | 5.95 | 10.42 | 1.04E-06 | hypothetical protein |
| TcSx10.chr2.007290_915550 | 5.95 | 10.42 | 1.04E-06 | hypothetical protein |
| TcSx10.chr2.007290_915800 | 5.95 | 10.42 | 1.04E-06 | hypothetical protein |
| TcSx10.chr4.021570_741300 | 4.45 | 9.00 | 5.39E-05 | microtubule-associated protein putative (fragment) |
| TcSx10.chr4.021570_748650 | 4.45 | 9.00 | 5.39E-05 | microtubule-associated protein putative (fragment) |
| TcSx10.chr4.021570_751350 | 4.45 | 9.00 | 5.39E-05 | microtubule-associated protein putative (fragment) |
| TcSx10.chr25.108900_519400 | 3.79 | 8.86 | 0.000359 | hypothetical protein |
| TcSx10.chr2.011350_1858158 | 7.41 | 8.64 | 3.21E-09 | ADP-ribosylation factor 1 putative |
| TcSx10.chr10.055590_1084550 | 3.59 | 8.59 | 0.000537 | hypothetical protein |
| TcSx10.chr10.055590_1086100 | 3.59 | 8.59 | 0.000537 | hypothetical protein |
| TcSx10.chr6.033680_631050 | 5.60 | 7.82 | 2.7E-07 | vacuolar protein sorting-associated protein-like putative |
| TcSx10.chr10.054250_760550 | 5.62 | 7.34 | 6.46E-08 | sarcoplasmic reticulum glycoprotein putative |
| TcSx10.chr7.035630_32700 | 4.37 | 7.11 | 4.35E-06 | transcription factor-like protein |
| TcSx10.chr30.127010_849009 | 2.75 | 6.78 | 0.001083 | vacuolar-type proton translocating pyrophosphatase 1 putative |
| TcSx10.chr7.039950_1096750 | 3.70 | 5.97 | 7.91E-07 | delta-1-pyrroline-5-carboxylate dehydrogenase putative |
| TcSx10.chr2.005050_510950 | 4.78 | 5.61 | 2.48E-11 | hypothetical protein |
| TcSx10.chr25.108680_467650 | 3.44 | 4.67 | 5.23E-10 | peptidyl-prolyl cis-trans isomerase putative |

## DISCUSSION

The development of vaccines for CD has been hindered by the lack of systematic studies to identify vaccine targets and an incomplete understanding of the mechanisms of parasite immune evasion and tissue persistence. This is further constrained by the diversity of parasite strains and their large number of multigene families encoding variant antigens (*14, 29*). In this work, we screened a *T. cruzi* genome-wide library for YSD using antibodies from CD patients or healthy donors and identified hundreds of *T. cruzi* proteins that are immunogenic in humans, determining their IRs. Using a combination of approaches, including quantitative proteomics of infectious forms, IR conservation among *T. cruzi* strains, and IR sequence divergence from human proteins, we prioritized 143 IRs as potential vaccine targets. Mice immunized with yeast expressing IR1 showed controlled infection in both acute and chronic murine CD, indicating the approach’s validity for identifying vaccine targets and yeast as a potential vehicle for vaccination. Notably, therapeutic vaccination of mice controlled parasite burden and led to the clearance of muscle tissues, indicating that a vaccine to treat chronic CD could be feasible.

The YSD-antibody screen revealed hundreds of IRs, several of which (∼23%) overlapped with peptides screened in previous studies using patients’ sera (*27, 28*). This overlap confirms the immunogenicity of some IRs, as shared sequences were recognized by the sera of dozens of patients. Because our *T. cruzi* genome-wide library is ∼30-fold the size of the parasite genome, with ∼270 clones/gene, and over 80% of the yeast express the library proteins on the surface, the entire parasite proteome is expected to be represented. This was corroborated by computational analysis, which indicated >10^5^ *T. cruzi* proteins or fragments expressed in frame, i.e., ∼18-fold the proteome. This number indicates the expression of many overlapping protein fragments, helping to map antigen-antibody binding sites. The robustness of the approach was also confirmed by the identification of several known immunogenic proteins, including those used in diagnostic tests for CD, e.g., surface antigen CA-2 (*43*), and studied for vaccines (*20, 21, 32, 37*). Several new CD-specific IRs (i.e., 918 immunogens) were identified that were not found in previous studies, suggesting conformational epitopes and post-translational modifications required for antibody interactions present in the YSD system but absent from synthetic peptides or phage libraries (*27, 43*). Yet some surface antigen post-translational modifications in yeast – extensive mannosylation – differ from those in *T. cruzi*, which is rich in α-galactosyl and sialic acid (*44–47*). Thus, some antibodies may still not have been captured by this approach.

Nonetheless, hundreds of IRs were identified that are potentially useful for developing vaccines, diagnostics, and biomarkers of cure or disease progression. The identification of IRs that induce antibodies in CD patients and are very conserved with host proteins indicates potential autoantigens associated with chronic CD, some of which have been reported (*48, 49*). These antibodies might increase levels during chronic CD, potentially causing tissue damage. Their early detection could serve as biomarkers, such as for cardiac pathologies.

To identify IRs as potential vaccine targets, we analyzed those conserved across all six DTUs and excluded multigene-family proteins. Single-cell RNA-seq data showed that only one or a few of the multigene family genes are expressed in each parasite (*50, 51*). Moreover, we showed that *T. cruzi* switches the expression of these antigens each time it exits an infected cell (*14*). Although these antigens are highly immunogenic, their variable expression within the parasite population may help them escape immune response. Furthermore, there is greater variability in these gene sequences across strains, suggesting that they recombine frequently and that a vaccine based on these antigens may allow parasites to escape the immune system. Although experimental vaccination with these antigens provides some protection against laboratory strains (*20, 21*), their efficacy may not be warranted in the field, where many different strains circulate. An important prerequisite we have considered for a vaccine candidate is the expression of the candidate antigens across the infectious stages: MTs, CTs and AMs. Priming the immune system with antigens expressed by all infectious forms ensures consistent host immune surveillance independent of the parasite form. Our TMT-MS data (*14*) quantitatively identified ∼4,000 antigens expressed in the infectious forms, which, although a large number, still represent <50% of the proteome. This allowed us to narrow 143 IRs expressed in all infectious forms. This number is likely to increase with additional proteomic studies, since RNA-seq suggests an additional 502 IRs shared across the infectious stages. Additional datasets, such as those expressed solely on the parasite surface, might help further guide the selection of vaccine candidates.

The IR1 candidate is likely encoded by a flagellar protein. It was identified in the flagellar proteome of *T. cruzi* (*31*), as it was the orthologue in *T. brucei* (*30*). Although a functional annotation is not available for this protein (assigned as a hypothetical protein), the encoded domains suggest that it, although divergent, might be related to adenylate kinase 7, which is found in cilia (*52*), which is evolutionarily related to the flagellum. The IR1 is encoded by 63 aa at the N-terminus of the protein, which, in the AlphaFold 3 model, protrudes from the protein surface. Vaccination of mice with IR1 in a prophylactic vaccine model reduced parasite burden in blood and tissues, indicating that IR1 can induce an effective immune response against *T. cruzi*. This response correlated with increased antibody levels and enhanced peripheral CD4^+^ and CD8^+^ T cell activation and function in immunized mice. The stimulation of CD8^+^ T cells by IR1 is relevant, as these cells function to clear infected cells in tissues. Although both yIR1 and rIR1 reduced infection in the blood, control of tissue parasitemia was more effective in mice immunized with yIR1. Similarly, therapeutic vaccination with yIR1 resulted in complete clearance of muscle tissue and a drastic reduction in intestinal tissue infection at 100 dpi, accompanied by increased IR1-antibody production and elevated CD8^+^ T cells. Both experiments suggest that yIR1 induces an effective immune response. The low/variable infection levels in other organs, e.g., adipose and heart, precluded accurate quantitative analysis. It is possible that the low parasite levels reflect dead parasites, as the quantification method relied on qPCR of parasite DNA, or tissue contamination during its resection. Alternatively, low parasite levels could be maintained in some tissues (e.g., adipose) due to an inefficient tissue-specific immune response. We also noted that parasite levels in tissues were often lower at 100 dpi than at 55 dpi. Despite the different vaccination approaches, the lower parasite levels at 100 dpi might reflect prolonged immune surveillance, as the timing of immune clearance likely differs between tissues. A longitudinal study would be required to determine whether these tissues could be cleared of parasites, since parasites that escape the tissues are likely cleared in the bloodstream. Infections with more virulent strains might help determine the efficacy of vaccination in these tissues, although highly virulent strains tend to be lethal to mice (*24*) and prevent analysis of chronic infections.

Although rIR1 and especially rIR1/Adj induced a significant level of peripheral CD8+ T cells in the prophylactic vaccination, the IFN-γ levels in mice did not increase proportionally. Exhaustion of CD8^+^ T cells has been reported in *T. cruzi* infection, leading to limited immune control of parasites in tissues (*39*). The apparently inefficient CD8^+^ T cells might have resulted from their over-stimulation by the combined effects of the high parasite load (during acute infection) and the adjuvant. In contrast, yIR1 vaccinations induced high antibody levels and a mild increase in peripheral CD4^+^ and CD8^+^ T cells compared to rIR1 and rIR1/Adj. This suggests that the mechanism of clearance may depend on elevated anti-IR1 antibody levels and a balanced cellular response, as high T cell counts may not necessarily confer better protection.

Downstream work measuring tissue-resident T cells might be necessary to elucidate their role in tissue protection. For example, the addition of IL-10/GM-CSF to yIR1 increased peripheral blood CD4^+^ and CD8^+^ T cells and IFN-γ levels during therapeutic vaccination, but did not confer greater protection than yIR1 alone. The lower levels of peripheral blood CD4^+^ and CD8^+^ T cells in yIR1-vaccinated mice may have also resulted from increased migration of these cells into target tissues. Markedly, vaccinations with yIR1 were more effective than those with rIR1 or its adjuvanted versions, indicating that yeast can serve as a vaccine vehicle in CD and likely also function as an adjuvant. In contrast, the IL-10/GM-CSF adjuvant added little to the yeast-based vaccine, and its beneficial effect might be associated with the choice of vaccine delivery, such as nucleic acids (*23, 36*).

Although prophylactic IR1 vaccinations resulted in controlled infection, they did not block infection. A blocking vaccine may depend on strong antibody responses elicited rapidly during acute infection. A further screen using sera of acutely infected CD patients might help identify antigens that induce rapidly produced antibodies. IgM antibodies are typically produced rapidly; however, they may not confer long-term protection, unlike IgG antibodies (*53*). Thus, a combination of targets to induce both types of responses: rapidly produced antibodies to block infection and long-lasting antibodies to clear persistent parasites, might be an alternative approach. The YSD screen was performed using sera from patients with chronic CD, which are biased toward antigens that elicit long-lasting antibody responses, a feature beneficial for a therapeutic vaccine, as observed in this work. Given the spread of CD into non-endemic areas and current estimates of ∼10.5 million infected individuals worldwide (*2*), a therapeutic vaccine is highly desirable, and yIR1 could be a promising candidate. A vaccine delivered via yeast benefits from high protein surface expression, proper protein folding and potential modifications, and yeast adjuvant properties. Moreover, yeast growth and heat inactivation/lyophilization are standard procedures, which could make production affordable for neglected diseases. Baker’s yeast allergies in the general population are extremely rare, and yeast advanced genetics could help eliminate allergens, e.g., enzymes and carbohydrates (*54*), to increase vaccine safety.

In summary, our *T. cruzi* antigen-antibody screen identified several immunogens for the development of diagnostics, biomarkers, and vaccines for CD. We validated yIR1 as a vaccine target and demonstrated potential efficacy for the therapeutic treatment of chronic infections. We also validated yeast as a vaccine vehicle that will likely accelerate vaccine discovery for CD and perhaps other parasitic infections.

## MATERIALS AND METHODS

### Parasite cell culture

Epimastigote cells of *Trypanosoma cruzi* strain SylvioX10 (ATCC 50823) were cultured in Liver Infusion Tryptose (LIT) medium (*14*), containing 10% fetal bovine serum and 1% penicillin-streptomycin. From a 12-day stationary culture of epimastigotes, 15 ml of cells were collected, and MTs were purified using DEAE-Sephadex resin. Metacyclic trypomastigotes were collected and washed with sterile PBS by centrifugation (1000 × g, 10 min at 4°C) for mouse infections. H9c2 cells were cultured in DMEM supplemented with 10% FBS at 37 °C in 5% CO2 (*14*).

### YSD library construction and protein expression

Genome-wide yeast surface display libraries were constructed from the *T. cruzi* Sylvio X10 strain genome (100% genome coverage) and from the *T. cruzi* CL Brenner Non-Esmeraldo-like strain genome (68% genome coverage). The libraries were mixed and transformed into EBY100 *Saccharomyces cerevisiae* yeast (ATCC MYA-4941) (*25, 26, 55*). The estimated number of cloned fragments required for genome coverage was determined using the Carbon and Clarke equation, as described (*25, 26*). To express *T. cruzi* proteins on the yeast surface, 1.2×10^7^ yeast cells were grown in YPD media (YPD – 1% yeast extract, 2% peptone, 2% dextrose, pH 6.8) at 30 °C and 225 rpm until OD_600_ 1. Cells were collected by centrifugation at 3000 *xg* for 5 min and washed three times in sterile MilliQ water by centrifugation at 3000 *xg* for 5 min. Cells were resuspended in 50 ml SD/-trp media (pH 5.8) containing 2% dextrose (SD/-trp+dex). Cells were incubated at 30 °C and shaking at 225 rpm until OD_600_ reached 1, approximately 16 h. Cells were collected by centrifugation and washed in water, as indicated above, followed by resuspension in SD/-trp with 2% raffinose (SD/-trp + raf) and incubating at 30 °C and 225 rpm for 2 h. Cells were collected by centrifugation and resuspended in SD/-trp + raf +1% galactose for protein expression and incubated at 30 °C and 225 rpm for 16 h. Yeasts were washed three times with sterile MilliQ by centrifugation at 3000 *xg* for 5 min.

### YSD antigen-antibody screen

The YSD screen was performed as described (*56*). Briefly, yeasts expressing the *T. cruzi* library growing in SD/-trp + raf +1% galactose for library protein expression were collected by centrifugation (3000 *xg* for 5 min), washed three times in ice-cold PBS by consecutive centrifugations (3000 *xg* for 5 min), and incubated in 1:1000 pooled sera of five chronic stage Chagas disease patients (diluted in PBS or 1:1000 pooled sera of two healthy individuals. Sera from CD patients from Bolivia or HI, aged 28-56 years, were kindly provided by Dr. Momar Ndao (McGill University Health Centre). Written informed consent was obtained from all participants. Serological tests were used to confirm the status of patients and control sera (*57*). Yeast and sera mix were incubated rotating for 2 hour at 4 °C. Cells were collected by centrifugation (3000 *xg* for 5 min) and washed three times in PBS by consecutive centrifugations (3000 *xg* for 5 min). Cells were resuspended in 400 µL 10 mM PBS and 100 µL magnetic Protein G beads and incubated at 4°C with rotation for 30 min. The 500 µL yeast and magnetic bead mixture was loaded onto a column (Miltenyi Biotec) and gravity-flowed through on a magnetic stand. Bound yeast were washed in the column three times in 5 mL PBS to remove nonspecific binders. Antibody-bound yeast was removed from the magnetic rack and re-cultured to expand the enriched (binders) population in SD/-trp +dex overnight. This process was repeated three times for both conditions to enrich for antibody-binding populations. After three rounds of enrichment, libraries were recovered from yeast and sequenced using Oxford Nanopore sequencing.

### Nanopore sequencing library preparation

Plasmid DNA was extracted using an adapted protocol from Sambrook & Russell (2001) (3). Pellets of yeast expressing *T. cruzi* YSD library (TcYSD) were resuspended in sorbitol buffer (1M sorbitol, 0.1M EDTA) with 12.5 mg/mL Zymolase and incubated at 37 °C for 1 h. Cells were collected by centrifugation at 15,000 *xg* for 1 min and resuspended in an equal volume of yeast resuspension buffer (50 mM Tris-HCl, 20 mM EDTA) and 10% SDS. Cells were incubated at 65 °C for 30 min, followed by the addition of 5M potassium acetate. The mixture was incubated on ice for 1 h, then centrifuged at 15,000 *xg* for 5 min, and the supernatant was collected. DNA was precipitated by adding an equal volume of isopropanol, followed by centrifugation at 15,000 *xg* for 15 min. The pellet was dissolved in TE buffer (10 mM Tris, 1 mM EDTA in MilliQ H2O, pH 7.2) and RNase A and incubated at 37 °C for 30 min. DNA was purified by adding 0.70x NucleoMag NGS beads (Takara Bio, Cat. #744970.5) to each sample and incubating for 15 min at RT. After being placed on a magnetic rack, the supernatant was removed, and the beads were washed two times via 70% EtOH and resuspended in TE to elute the DNA. To prepare samples for Oxford Nanopore Sequencing, *T. cruzi* gene fragments were amplified using primers 1 and 2 (**Table S5**), and the library was prepared as described (*26*).

### Screen computational analysis and IR cross-validation

All scripts used in this work are available on the Cestari Lab GitHub (https://github.com/cestari-lab). Nanopore sequencing fast5 data from sequenced libraries were base called with Guppy (Oxford Nanopore Technologies). Fastq sequences were mapped to *T. cruzi* Sylvio X10 strain genome using minimap2 (*58*). Mapping statistics and format conversions were obtained using Samtools (*59*). Samtools was used to remove secondary and supplementary alignments and to filter for MapQ> 1. Read counts per gene were obtained using featureCounts from Subread (*60*). The coverage was analyzed using plotCoverage from deepTools (*61*). The resulting .bed file was visualized using the Integrative Genomic Visualizer (IGV) (*62*), and plotted on a circular plot using the circus package (*63*). The estimated amino acid length was calculated using the libframe tool (*25*). Enrichment analysis of genes was performed using the EdgeR (*64*). To identify enriched immunogenic regions (IRs), the macs3 tool was used (*65*), using narrow Peaks and a q-value cut-off of 0.05. Using the genomic coordinates of IRs, the nucleotide FASTA files and protein FASTA files were obtained in the correct frame using an in-house Python script: extract_regions.py https://github.com/cestari-lab/YSD). The IDs of genes encoding IRs were searched in TMT-MS data and RNA-seq data, the latter mapped to the *T. cruzi* Sylvio X10 genome (*14*) to identify IRs expressed in infectious forms of *T. cruzi*. To identify peptides that react with CD patient antibodies matching to IRs, we collected 3868 peptide sequences that react with 71 CD patients’ sera, i.e., peptides from the Epitope Atlas (*27*) as well as peptide sequences from the Immune Epitope Database (IEDB) using the search parameters: Epitope: ‘Linear peptide’; Assay: only ‘B Cell’ and ‘Positive’; Organism: ‘Trypanosoma cruzi (ID:5693)’; MHC Restriction: ‘Any’; Host: ‘Human’; Disease: ‘Any’.

Peptide sequences from previously identified peptides were compared to IRs using the blastp tool. Matches were filtered using a 100% identity threshold and an e-value ≤ 0.05. To analyze IR sequences against different *T. cruzi* DTUs, we analyzed IR sequences using the blastn tool with an identity threshold of 80% and an e-value ≤ 0.05. IRs were compared against the genomes of *T. cruzi* Sylvio X10 (DTU I), *T. cruzi* 231 (Release 67), *T. cruzi* Bug2148 (Release 67), *T. cruzi* YC6 (Release 67), *T. cruzi* CL Brener Non-Esmeraldo-like (Release 67), and *T. cruzi* CL Brenner (Release 67) retrieved from TriTrypDB (*66*). IRs were compared to the human proteome using the UniProt ProteomeID: UP000005640 by blastp. Sequences with a percentage identity > 30 were discarded. A fasta file of all multi-gene family proteins (trans-sialidases, mucins, mucin-associated surface proteins, GP63, retro-transposon hotspots, dispersed gene family 1 (*14*)) was compared to the amino acid sequences of IRs filtered with an identity threshold of 100 and an e-value ≤ 0.05.

### Western blot

Protein extracts were prepared from 25 mL of yeast cells with surface protein expression (see TcYSD surface protein expression). Cells were harvested at 3000 *xg* for 5 min and resuspended in yeast lysis buffer (5 mM DTT, 1X proteinase inhibitor cocktail [Roche] supplemented with 5 mM phenylmethylsulfonyl fluoride [PMSF]), vortexed for 10 seconds, and rotated at 280 rpm at room temperature (RT) for 20 min. The mixture was centrifuged for 5 min at 13000 *xg*, and the supernatant was collected for Western blot analysis. 30 µL of protein lysate were mixed with 10 µL of 4x Laemmli buffer (0.25M Tris; 8% sodium dodecyl sulfate, 40% glycerol, 20% 2-mercapto-ethanol, 6 mM bromophenol blue) and incubate at 95 °C for 5 min.

Proteins were resolved in 12% SDS-PAGE and transferred to polyvinylidene fluoride (PVDF) membranes. Membranes were blocked with 6% milk in PBS-T(1X PBS, 0.05% Tween) for 1 hour at RT. Afterwards, membranes were incubated with either mouse α-Xpress antibodies (Life Technologies) diluted in 1:000 blocking buffer (6% milk in PBS-T), rabbit α-glycosome diluted in 1:5,000 blocking buffer, or rabbit α-Pex14 diluted in 1:5000 blocking buffer (Pex-14 and glycosome antibodies were kindly provided by Dr. Marilyn Parsons, Seattle Children’s) (*67*).

Membranes were incubated with primary antibodies for 2 h, then washed with 10 mL PBS-T. Membranes were then incubated with 1:5000 goat α-mouse IgG-HRP (Bio-Rad) in blocking buffer, or 1:5000 donkey α-rabbit IgG-HRP (Bio-Rad) in blocking buffer for 1 hour at RT. Membranes were then washed as above and then developed with ECL chemiluminescence solution (Life Technologies) using a ChemiDoc MP Imaging System and Image Lab software (Bio-Rad).

### Flow cytometry and fluorescence microscopy

Yeast cells (2×10^6^) were collected from culture by centrifugation at 4000 *xg* for 5 min. Media was removed, and cells were resuspended in 1 mL PBS and centrifuged at 4000 *xg* for 5 min. The procedure was repeated three times to wash the cells. Cells were fixed in 1 mL of 1% paraformaldehyde in PBS for 10 min and washed again as described above. Surface proteins were blocked with 1% bovine serum albumin (BSA) in PBS for 2 h and then collected by centrifugation at 4000 *xg* for 5 min. The cells were incubated in 1 mL of 1:500 mouse α-Xpress (Life Technologies) in 1% BSA in PBS for 2h. After incubation, cells were washed three times with PBS at 4000 *xg* for 5 min. Cells were then resuspended in 1 mL of 1:1000 goat anti-mouse IgG-AF488 (Life Technologies) in 1% BSA in PBS and incubated for 1 h. Cells were again washed three times with PBS at 4000 *xg* for 5 min and resuspended in 1 mL PBS. An aliquot of about 1×10^5^ cells was placed on a 24-well plate, covered with a coverslip and imaged using the Cytation 5 imaging platform using the BioTek Gen5 Software (Agilent BioTek). The remaining cells samples were acquired via Attune^TM^ NxT Acoustic Focusing Cytometer (ThermoFisher Scientific) using the Attune^TM^ Cytometric Software and samples were analyzed using FlowJo v10 Software.

### Cloning IRs for bacterial and yeast expression

Five IRs (see **Fig. 2I** legend for gene IDs) were amplified from *T. cruzi* Sylvio X10 genomic DNA using primer #3-12 (**Table S5**) for expression in yeast. *T. cruzi* Sylvio X10 genomic DNA was isolated using the Monarch Genomic DNA extraction kit (New England Biolabs), according to the manufacturer’s instructions. IR1 to IR5 PCR sequences were cloned into the pYD1 expression. Primers were designed for Gibson Assembly containing a BamHI restriction site, and the pYD1 vector was digested with BamHI. PCR was performed using Taq DNA Polymerase under the following conditions: 94 °C for 30 s, followed by 35 cycles of 94 °C for 30 s, 50 °C for 30 s, and 68 °C for 90 s, then a final extension at 68 °C for 5 min. For cloning IR1 into the pET28a(+) expression vector for protein expression and purification from *Escherichia coli*, the vector was digested with NdeI and XhoI. Primers (13–14) were designed for Gibson assembly (**Table S5)**. The purified PCR product and digested pET28a(+) or pYD1 vector were ligated at a 20:1 insert-to-vector molar ratio using HiFi Assembly Reaction Mix for at 50°C for 1 h. The ligation mixture was transformed into chemically competent *E. coli* DH5α cells via heat shock at 42 °C for 30 s, followed by recovery in SOC medium (2% tryptone, 0.5% yeast extract, 10 mM NaCl, 10 mM MgCl2, 20 mM glucose) at 37°C for 1 h. The IR1-containing pET28(+) vector was transformed into protein-expressing chemically competent *E. coli* NiCo21(DE3) via heat shock at 42°C for 30 s, followed by recovery in SOC medium at 37°C for 1 h. Transformants were selected on LB agar plates containing 50 µg/mL kanamycin. The IR1 containing pYD1 vector was transformed into EBY100 *S. cerevisiae*, as described (*56*).

### Expression and purification of rIR1

pET28a-IR1-transformed *E. coli* NiCo21(DE3) were cultured overnight in 80 mL of LB medium (10 g tryptone, 5 g yeast extract, 10 g NaCl, pH 7.3) containing 50 µg/mL kanamycin at 37°C and 225 rpm. The overnight culture was then diluted 50x in 4 L LB media containing 50 µg/mL kanamycin and grown at 37 °C and 225 rpm until OD_600_ reached 0.4 – 0.6. Once the desired OD_600_ was achieved, 0.2 mM isopropyl β-D-1-thiogalactopyranoside (IPTG) was added to induce protein expression, followed by a 4 h incubation at 37 °C with shaking at 225 rpm.

After protein induction, cells were harvested by centrifugation at 5000 *xg* for 30 min. Cells were then resuspended in 10 mL lysis buffer (10 mM PMSF, 2 mg lysozyme, 5% glycerol, 5 mM DTT in PBS) and sonicated at 1000 J, for 3 min (10 s ON; 50 s OFF) at 25% power. The lysate was clarified by centrifugation at 13000 *xg* for 5 min at 4 °C, and the supernatant was collected. Ni-NTA beads (Bio-Rad) were equilibrated in PBS pH 8.0 and added to the supernatant. The mixture was incubated with the beads at 4 °C for 2 h, rotating at 60 rpm. The resin was washed with 50 mL wash buffer (50 mM Tris-HCl, pH 8.0, 300 mM NaCl, 10 mM imidazole). Bound proteins were eluted using elution buffer (50 mM Tris-HCl, pH 8.0, 300 mM NaCl, 250 mM imidazole). Purified fractions were resolved in a 15% SDS-PAGE and analyzed by Coomassie staining. Confirmation of correct protein expression was determined by Western blot via 1:2000 mouse α-His-tag primary antibody (Invitrogen) and mass spectrometry of excised bands from 10% SDS/PAGE Coomassie stained (*68*) at the MUHC Proteomics Core. Pooled fractions were dialyzed in PBS and stored at -80 °C.

### Expression of IR1 in yeast and lyophilization

Yeast cells expressing IR1 were generated by transformation of EBY100 with the pYD1-IR1 vector, as described above in “YSD library construction and protein expression”. Following protein expression, 1×10^9^ yeast cells were washed three times with sterile MilliQ water at 3000 *xg* for 5 min. Yeast cells (1×10^7^ aliquots) were heat-inactivated at 60 °C for 90 min. Heat-inactivated yeast were lyophilized overnight using the Labconco Lyph-Lock 6 lyophilizer (Labconco), according to the manufacturer’s instructions. Afterwards, lyophilized yeast was kept at -80 °C until use. Before use, yeast cells were rehydrated in 1 mL sterile PBS.

### Immunizations and parasite challenge

Groups of six-to-ten-week-old BALB/c male mice (n = 5 per group) were purchased from Charles River Laboratories and maintained at the small animal research unit of McGill University. All animal procedures were approved by the McGill University animal committee, under protocol MCGL-10152. Mice were immunized intraperitonially (ip) with rIR1 (10 µg/dose/mouse) in PBS, alone or with recombinant IL-12 (1 µg/dose/mouse) and GM-CSF (50 ng/dose/mouse). Additionally, mice were immunized with yIR1 (1×10^6^ cells/dose/mouse) alone or in combination with rIL-10 and rGM-CSF as above. Control mice received adjuvant alone, or yeast (EBY100) expressing the empty pYD1 plasmid (1×10^6^ cells/dose/mouse). All injections were delivered in 100 µL of PBS. A naive group (non-immunized or infected) was also included and handled like the experimental groups. For prophylactic immunization, mice received a prime immunization followed by two booster immunizations at 14-day intervals. All groups were challenged ip infections with 1×10^6^ *T. cruzi* Sylvio X10 MTs two weeks after the last immunization. 5 µl of blood was collected via tail-prick every two days starting at 10 dpi. For therapeutic immunization, mice were infected with 1×10^6^ *T. cruzi* Sylvio X10 MTs; blood was collected via tail prick (10 µl) at 5 dpi and 20 dpi to confirm infection, and immunization treatments were initiated at 55 dpi (prime), followed by two booster immunizations at 7-day intervals. Blood was also collected at the endpoints (55 dpi for prophylactic immunization; 100 dpi for therapeutic immunization) via cardiac puncture (∼500 µl). Blood was coagulated at RT for 30 min before centrifugation at 1000 *xg* for 10 min to collect the serum. Serum was stored at -80 °C until use. Organs (large intestine, heart, adipose tissue, and hindlimb skeletal muscle) were also harvested. Animals were humanely euthanized via CO_2_ overdose and cervical dislocation at the experimental endpoint.

### Quantitative real-time PCR (qPCR) of blood and tissue parasite burden

Mice tissues were lysed, and gDNA was extracted from blood or 20-30 mg tissues using the Monarch^®^ Spin gDNA Extraction Kit (New England Biolabs). Samples were purified using NucleoMag NGS Beads (Takara Bio) at a 1:1 ratio and eluted in 40 µl sterile MilliQ water, 5 µl of purified sample were used per reaction. Amplification of *T. cruzi* satellite DNA was done using the specific primers cruzi1 and cruzi2 (*69*), both at 750 nM, and the TaqMan probe cruzi3 at 50 nM (**Table S5**). Samples were spiked with the tonoplast intrinsic protein 5;1 (TIP5;1; accession number: NM_114612) from *Arabidopsis thaliana* as an internal amplification control (IAC) (*69*). The IAC was amplified using primers IAC-F and IAC-R, each at 200 nM, and the IAC probe at 50 nM (**Table S5)**. All samples were run using the Luna Universal qPCR MasterMix (New England Biolabs). Standard curves were done by spiking naïve tissues (blood, large intestine, heart, adipose, and skeletal muscle) with 10-fold serial dilutions of 10^6^ parasites/mL to 1 parasite/mL. PCR conditions consisted of holding at 50 °C for 2 min, 95 °C for 10 min, followed by 40 cycles of 95 °C for 15 seconds and 58 °C for 1 minute. Each sample was run in duplicate, and fluorescence was collected after each cycle on the Step One Plus Real Time PCR System (Applied Biosystems). For tissues extracted at the endpoint, samples were normalized using murine GAPDH amplification (*70*). Briefly, 5 µl of purified gDNA extracted from tissue was mixed with 250 nM of GAPDH-F and 250 nM of GAPDH-R primers (**Table S5**). Samples were added to the BlasTaq™ 2X qPCR MasterMix (Applied Biological Materials). PCR conditions consisted of a hold at 95 °C for 20 seconds, followed by 40 cycles of 95 °C for 1 second and 60 °C for 10 seconds. Each sample was run in duplicate, and fluorescence signals were collected after each cycle using the StepOne Plus Real-Time PCR System (Applied Biosystems).

### Reverse-transcription quantitative PCR (RT-qPCR) of cytokines from blood

At the experimental endpoint, 50 µl of blood was collected and added to 500 µl of TRIzol (ThermoFisher Scientific). To each sample, 100 µl of phenol:chloroform:isoamyl alcohol (25:24:1) (BioShop) was added and mixed by vortexing, followed by centrifugation at 10,000 x*g* for 10 min; the clear phase containing RNA was collected. To the clear phase, 0.5x 100% EtOH was added and mixed by vortexing. Samples were placed on the columns of the 96-Well Plate Bacterial Total RNA Miniprep Super Kit (BioBasic), and RNA was collected from the columns by centrifugation at 2000 *xg* for 5 min. Samples were washed with 500 µl of Universal GT buffer and centrifugation at 2000 *xg* for 5 min, followed by 500 µl of Universal NT buffer and centrifugation at 2000 *xg*. RNA was collected by adding 50 µl of RNase-free water and then centrifuged at 2000 *xg.* From the RNA, cDNA was synthesized by first adding 5 µl total RNA, 2 µl 60 µM random primer mix (Invitrogen), 1 µl of 10 mM dNTPs, and 2 µl of RNase-free water. This reaction was incubated at 65 °C for 5 min. After incubation, samples were immediately placed on ice, and 2 µl of M-MuLV buffer (New England Biolabs), 1 µl M-MuLV Reverse Transcriptase (New England Biolabs), 0.2 µl RNase inhibitor, and 6.8 µl RNase-free water were added to each sample. Samples were incubated at 25 °C for 5 min, followed by 42 °C for 90 min, and 65 °C for 5 min. cDNA was diluted 1:10 prior to RT-qPCR in water. Briefly, 5 µl of cDNA extracted from blood was mixed with 250 nM of forward and reverse primers for IFN-γ, TGF-β, IL-4, and IL-10 (**Table S5)**. Additionally, cDNA was mixed with 250 nM of forward and reverse primers against GAPDH and actin for relative quantification (**Table S5**), using cDNA without reverse transcription as a control for DNA contamination. Samples were added to the BlasTaq™ 2X qPCR MasterMix (Applied Biological Materials). PCR conditions consisted of a hold at 95 °C for 20 seconds, followed by 40 cycles of 95 °C for 1 second and 60 °C for 10 seconds. Each sample was run in duplicate, and fluorescence signals were collected after each cycle using the StepOne Plus Real-Time PCR System (Applied Biosystems).

### Enzyme-Linked Immunosorbent Assays (ELISA)

100 ng of rIR1 was incubated in 100 µl of carbonate coating buffer (0.1 M Na_2_CO_3_, 0.1 M NaHCO_3_, pH 7.5) overnight at 4°C on ELISA High-Binding 96-well plates (Sarstedt). Wells were washed 3 times with 200 µl of PBS and blocked for 2 h with 100 µl of 3% soy trypticase broth in PBS (STBP). Samples were added in duplicate to the wells at a 1:500 dilution in 3% STBP and incubated for 2 hour at RT. Wells were washed three times with 200 µl PBS. Samples were incubated for 1 h with 100 µl of a 1:4000 dilution of goat anti-mouse IgG-HRP-conjugated secondary antibody (Bio-Rad) in STBP. Wells were washed as indicated above. Samples were quantified by adding 150 µl of the 1-Step™ ABTS Substrate Solution (ThermoFisher Scientific), and the optical density was measured at 415 nm using the BioTek Synergy H1 Micro-Plate Reader (Bio-Rad). Cytokine expression was evaluated following the Mouse IFN-gamma ELISA Kit (ABclonal) and the Mouse IL-10 ELISA Kit (ABclonal). Mouse serum was diluted 1:100 in PBS to evaluate IFN-γ and 1:20 in PBS to evaluate IL-10. Optical density was measured at 450 nm using the BioTek Synergy H1 Micro-Plate Reader (Bio-Rad).

### T cell analysis by flow cytometry

Peripheral blood mononuclear cells (PBMCs) were prepared by retrieving 50 µl of blood via cardiac puncture at the experimental endpoint, mixed with 500 µl Ammonium-Chloride-Potassium (ACK) lysis buffer (150 mM NH_4_Cl, 10 mM KHCO_3_, 0.1 mM Na_2_EDTA, pH 7.2), and incubated on ice for 5 min to lyse red blood cells. The lysis was quenched by adding 500 µl of PBS buffer, followed by centrifugation at 350 *xg* and 4 °C for 15 min. The supernatant was removed, the cell pellet resuspended, and washed in 1 ml of PBS, and the cells were centrifuged at 350 *xg* and 4 °C for 15 min. Cells were resuspended in 200 µl PBS, and 2×10^6^ cells/well were plated on a V-bottom 96-well plate (Ultident Scientific). Cells were collected at 1500 *xg* for 5 min at 4 °C, resuspended in 50 µl of 1:1000 eBioscience™ Fixable Viability Dye eFluor™ 506 (Thermo Fisher Scientific), and incubated at 4 °C for 15 min, protected from light. To wash cells, 150 µl of BD Pharmingen™ Stain Buffer (FBS) was added to wells and cells were collected via centrifugation at 1500 *xg* for 5 min at 4 °C. The wash was repeated by resuspending cells in 200 µl of FBS and centrifuging as indicated above. The cells were resuspended in 50 µl of BD Pharmingen™ Purified Rat Anti-Mouse CD16/CD32 (Mouse BD Fc Block™) diluted to 1:100 and incubated in the dark for 10 min at RT. Surface antibodies were added after 10 min without washing. The antibodies BD Pharmingen™ FITC Hamster Anti-Mouse CD3e, BD Pharmingen™ APC-Cy™7 Rat Anti-Mouse CD4, BD Horizon™ V450 Rat anti-Mouse CD8a, BD Pharmingen™ APC Rat Anti-Mouse CD44, BD Horizon™ BV711 Rat Anti-Mouse CD62L were diluted 1:400 in 100 µl of PBS. The cells were incubated at 4 °C for 30 min. Cells were washed 3 times in 200 µl of FACS buffer and collected at 1500 x g for 5 min. Samples were resuspended in 200 µl FACS buffer, and acquired via Attune^TM^ NxT Acoustic Focusing Cytometer (ThermoFisher Scientific) using the Attune^TM^ Cytometric Software. Data were analyzed using FlowJo v10 Software. Gates were set for live cells (eBioscience™ Fixable Viability Dye eFluor™ 506 labelling), and then for T cells (FITC-CD3e labelling), using forward and side scatter properties. The percentage of activated CD4^+^ and CD8^+^ T-cells was determined based on gated CD3^+^ T-cells (**Figure S7**).

### Statistical analysis

Statistical analysis was performed in RStudio (*71*). All experiments were performed with at least three biological replicates. Results were considered statistically significant at α = 0.05. All reported confidence intervals are at the 95% level. Statistical analysis of library enrichment was performed using the edgeR quasi-likelihood pipeline (*64*). Genes were considered differentially enriched with a fold-change ≥ 2 and a false discovery rate (FDR)-adjusted *p*-value ≤ 0.05. For vaccination experiments, data were evaluated for normality and homogeneity of variance using the Shapiro-Wilk (and Quantile-Quantile plot) and Levene’s tests, respectively. Normally distributed, homoscedastic data were analyzed using ANOVA followed by Tukey’s Honest Significant Difference (HSD) post-hoc test. Non-parametric or heteroscedastic data were analyzed using non-parametric multiple contrast test procedures (MCTP) with Dunnett-type contrasts to compare experimental groups against the control group.

## List of Supplementary Materials

Figure S1 to Figure S8

Table S1 to Table S7

Data file S1 to Data file S4

## Acknowledgments

This research was partly enabled by computational resources provided by Calcul Quebec (https://www.calculquebec.ca/en/) and the Digital Research Alliance of Canada (alliancecan.ca). We thank Dr. Ephraim Ansa-Addo (Pelotonia Institute for Immuno-Oncology, The Ohio State University, USA) for sharing protocols and advice on T-cell analysis, reading and providing valuable comments to the manuscript.

## Author contributions

Conceptualization: ML, IC

Methodology: ML, LCS, IC.

Investigation: ML, LCS, VBA, IC

Visualization: ML, IC

Funding acquisition: ML, IC

Project administration: IC

Supervision: IC

Writing – original draft: ML, IC

Writing – review & editing: ML, LCS, VBA, IC

## Data and materials availability

Computer codes used in this manuscript are available at GitHub: https://github.com/cestari-lab/YSD. Oxford Nanopore Sequencing (fastq) are available in the Sequence Read Archive (SRA) with the BioProject identification PRJNA1477972 (https://www.ncbi.nlm.nih.gov/bioproject/1477972). Plasmids and cell lines are available upon request and may require completion of a materials transfer agreement. All data are available in the main text or the supplementary materials.

