## Supplementary Information for "A genome wide antigen-antibody screen identifies a yeast-based therapeutic vaccine candidate for Chagas disease"

Mira Loock[1], Lissa Cruz-Saavedra[1], Valeria Bernal Araujo[1], Igor Cestari[1*]
[1]Institute of Parasitology, McGill University; Sainte-Anne-de-Bellevue, QC H9X 3V9, Canada.

### List of Supplementary Materials

Figure S1 to Figure S8
Table S1 to Table S7
Description of Data file S1 to Data file S4

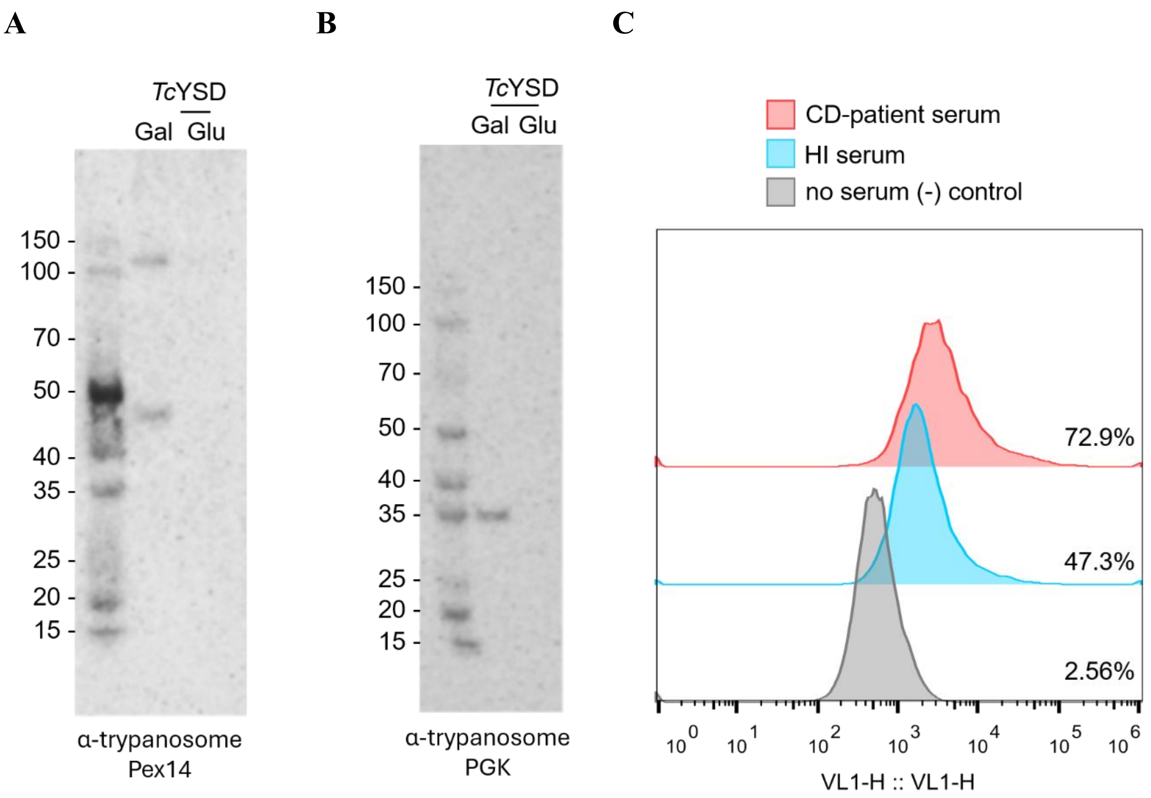


**Figure S1. YSD expression of *T. cruzi* proteins. A-B)** Western blot of lysate from yeast expressing the *T. cruzi* genome-wide library (*Tc*YSD). Expression of trypanosomes peroxin 14 (Pex14) and phosphoglycerate kinase (PGK) proteins was detected with specific antibodies (see Table S7). α-trypanosome PGK recognizes PGKA, B, and C. Yeast was induced to express the *T. cruzi* library with 1% galactose (Gal), or repressed with 1% glucose (Glu, i.e., no expression). The protein migration size reflects the size of the protein fragments expressed in the library fused to Aga2p. **C)** Flow cytometry analysis of the *Tc*YSD (induced with 1% galactose) binding to antibodies from Chagas disease (CD) patients' pooled serum or healthy individuals' (HI) pooled serum. Sera were diluted 1:1000 and probed with goat anti-Human IgG Alexa Fluor Plus 405.

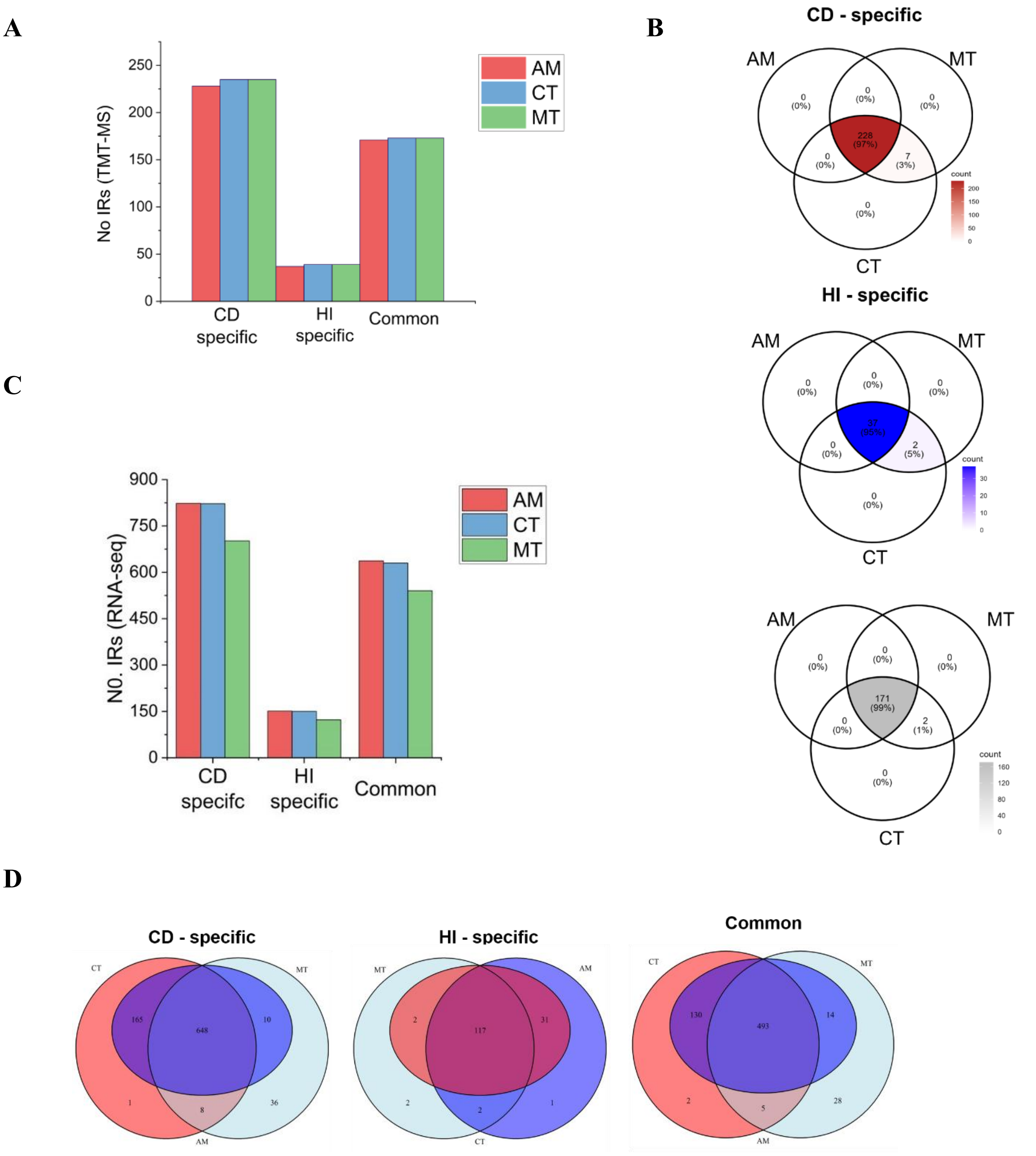


**Figure S2. Analysis of IR expression by proteomics and RNA-seq. A)** Counts of IRs identified in the proteomic dataset at each infectious life stage. AM, amastigote; MT, metacyclic trypomastigote; CT, cell-derived trypomastigote. No, number. TMT-MS, Tandem mass-tag mass spectrometry. **B)** Venn diagram of IRs found at each life stage (AM, MT, CT), for CD-specific IRs, HI-specific IRs, and IRs common between datasets. **C)** RNA-seq identification of IRs at each infectious life stage (AM, MT, CT) from CD-enriched, HI-enriched, and common IRs. **D)** Venn diagram of IRs found at each life stage (AM, MT, CT) from RNA-seq dataset.

A

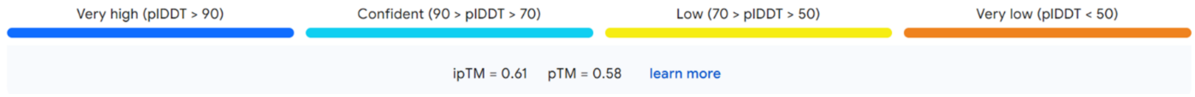


B

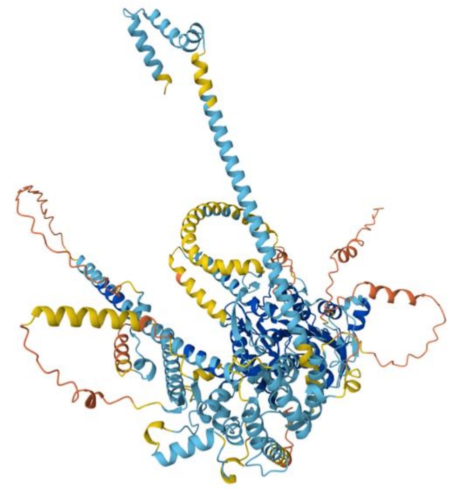

C

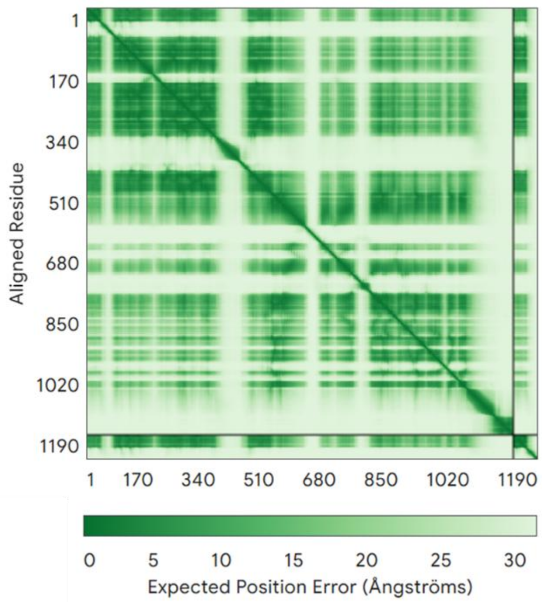


D

**>TcSx10.chr5.027000.m01.polypeptide**
ID=TcSx10.chr5.027000.m01.polypeptide|Name=TcSx10.chr5.027000.m01.polypeptide|organism=Trypanosoma trypanosoma cruzi|type=polypeptide|length=1128|location=TcSylvioHIFI_5:166962..170348+

**MKRVLVLNVDSFVGREVCRCFFDAKEYLVDGTLYHFQSTQGSPEARESLRTDGLDSAGTIKPH**CATASTSSVRLDVPPDMRPFLNS
VMPRHGDISDEQFRVRVLTYDIIIAILEEDAFEADCTIKVLRGTHYEVEKTFVLVSNVFTWTQTDAHERELRRAARRAEREAARAQRDED
DVIEEEEEEEEEEEEEEMEQKVWTEEDYRQRYPDLRYHLWKDLEHAVKHANSETLHTYVLWAGLPYGRGEDLLASHFNAAWRHQEML
QYGDGSNYIPTIHVKDLARIIYLVGSSYDTLEDRYMFAVDQGNNTQSDILQGIKDFIGGAVSITAPEIDRKNQVVIPDNQKKDRREADVKE
NELAEEIQHEHDVLVELGMRESQVLDEAKEEMLRIKEISQREKLLRLGLGIQEIENDDDTNHALSPEAASKVGASLTIAPPHGLANWFSA
VDMRCEPGAVLALLEEEEWVSLGGFFANVDKIVQEFKAARQLRPMRLVLSGPPLSKVGEVAGVLAELFDIPHLTLANVTAAYEEHVRSL
REELIDILVLRQLRRHTAREERRRREKERAKDARRKAREEEGEEGEEGEEEEEEEEEEEGEEEGEEDEEEGAEEEEEEKGEEGTPAAQDLDP
AERIRAAITEAMLLHAPEDEEDEEEDDSHYNADDEEEVDEKAAAMAQQVDEAERRRIATLREEYRFATQLLSLRLLNGEFPRRPKSADE
EEEGEEEEEEYDEYDDEEYMNRKKRREQKKDAKKDVTEEPPPTIRYFDEVMAVMVRWRLRQEDCKNQGYILEAFPETVRQAWLTFLAD
TEEQKEMQRPKLQRVRDEEAEPQPLLDANFPLPVLADLEEDRKQEIMELMTGRRFASAEEVLVPTHDAFFPDCFIALTGNEVTLRSQIYA
AIATATPGDTTPDFSEQFTRYMTHNRPGAPPTESLVCWFQTVVAERVDEDGDVAVATEGGELREKVPRKACVLFVPVFSTFPQAVLPKFFN
REEISGMKAYCTFIRSEILKNIGLDSCGSVIDSMRAEAEANKSRIQELASEREKLEKLSASDKLFAAAAQKLEDDFLENERRALRFYQAKE
ELASIVGIDSIPVDVYLMRYVMPSLTPLMAEVVRMRPEDPVTVLADALFHHKRQVSL

**Figure S3. Predicted AlphaFold 3 structure of the protein encoded by TcSx10.chr5.027000.** **A)** Shows colour-coded predicted local distance difference test (pLDDT) indicating local quality of the folding of different protein segments; predicted template modelling (TM)-score (pTM) indicating the global structural accuracy on a scale of 0-1; and interface pTM indicating confidence in the interactions of different chains or domains on a scale from 0.1. **B)** Colour-coded protein structure. Colours match the pLDDT graph in A. **C)** Predicted aligned error (PAE) plot showing confidence in the relative positions of different parts of the protein. **D)** Predicted amino acid sequence of the gene encoding TcSx10.chr5.027000. IR1 sequence is shown in bold.

**A**

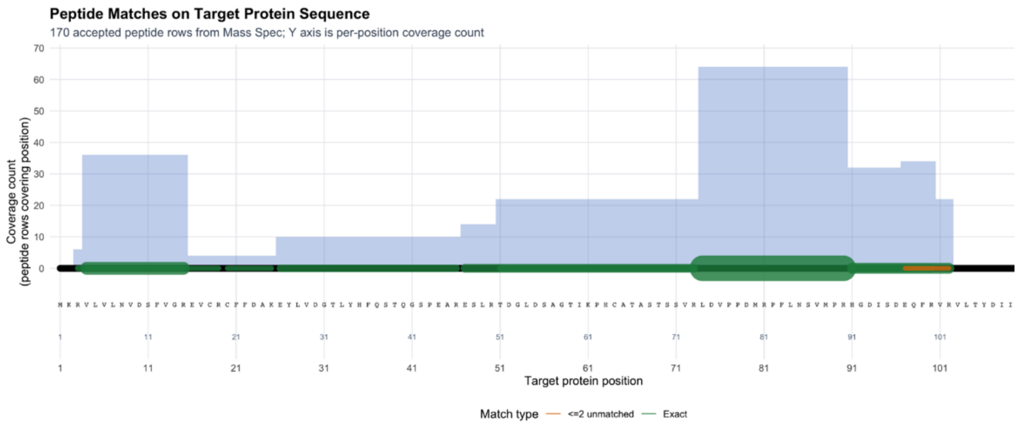


**B**

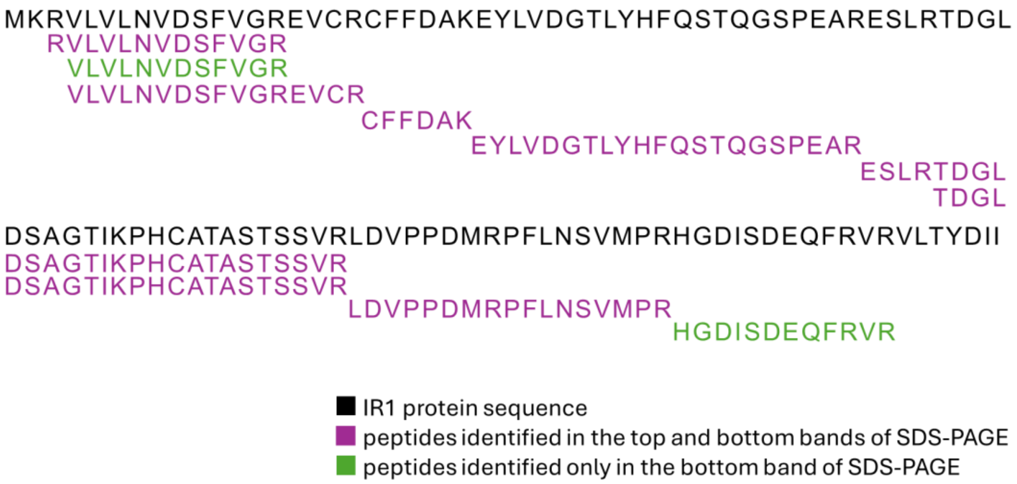


**Figure S4. Identification or rRI1-His by mass spectrometry. A)** Peptide coverage over the 116 aa of the recombinantly expressed His-tagged IR1-encoding protein (rIR1) identified by mass spectrometry of the bands excised from 15% Coomassie-stained SDS/PAGE (as in Fig. 5B). **B)** Unique peptide sequences matching the sequence of the expressed protein identified by mass spectrometry. The colour indicates peptides found in the bottom and top bands (bottom band is the predicted 12.7 KDa molecular weight of rIR1), and the top band oligomeric forms observed in SDS/PAGE and Western blot (as in **Fig. 5B**).

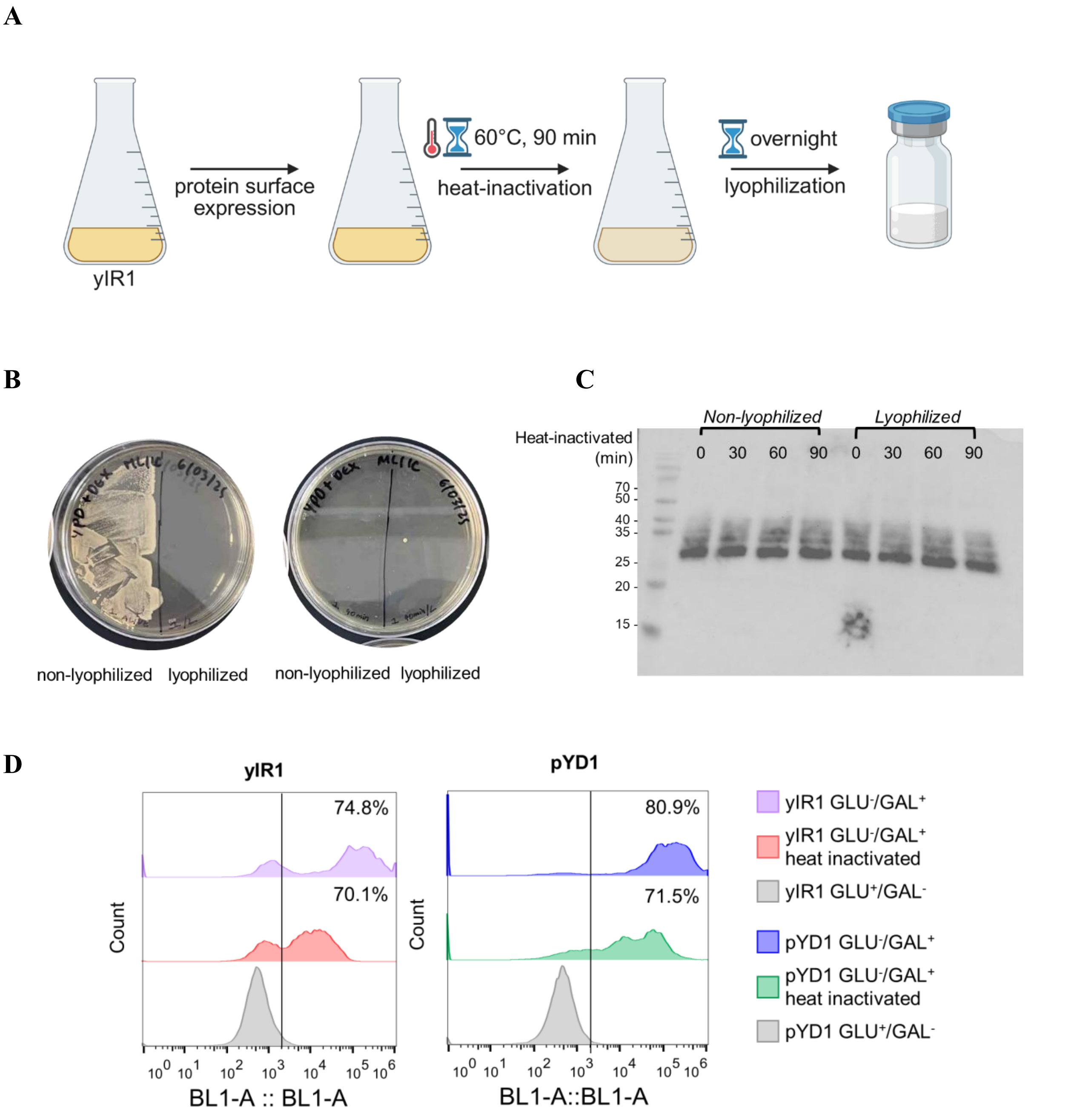


**Figure S5. Preparation of yIR1 for vaccination. A)** Diagram of the steps taken from yeast expression to heat inactivation and lyophilization. Created in BioRender. Cestari, I. (2026) https://BioRender.com/atymc6d. **B)** YPD/Agar plate of yIR1 before (0 min) and after 90 heat-inactivation at 60 °C (90 min), and before and after lyophilization. **C)** Western blot of yIR1 lysate before and after lyophilization and heat inactivation at multiple time points. Western blotting was developed with α-Xpress antibodies. **D)** Flow cytometry of yIR1 or yeast expressing pYD1 (empty vector, results in expression of Aga2p without IR1) with α-Xpress antibodies and α-mouse IgG-Alexa Fluor488 before and after heat inactivation. Non-induced yeast (yIR1 Glu+/Gal- or pYD1 Glu+/Gal-) was included as a control.

**A**

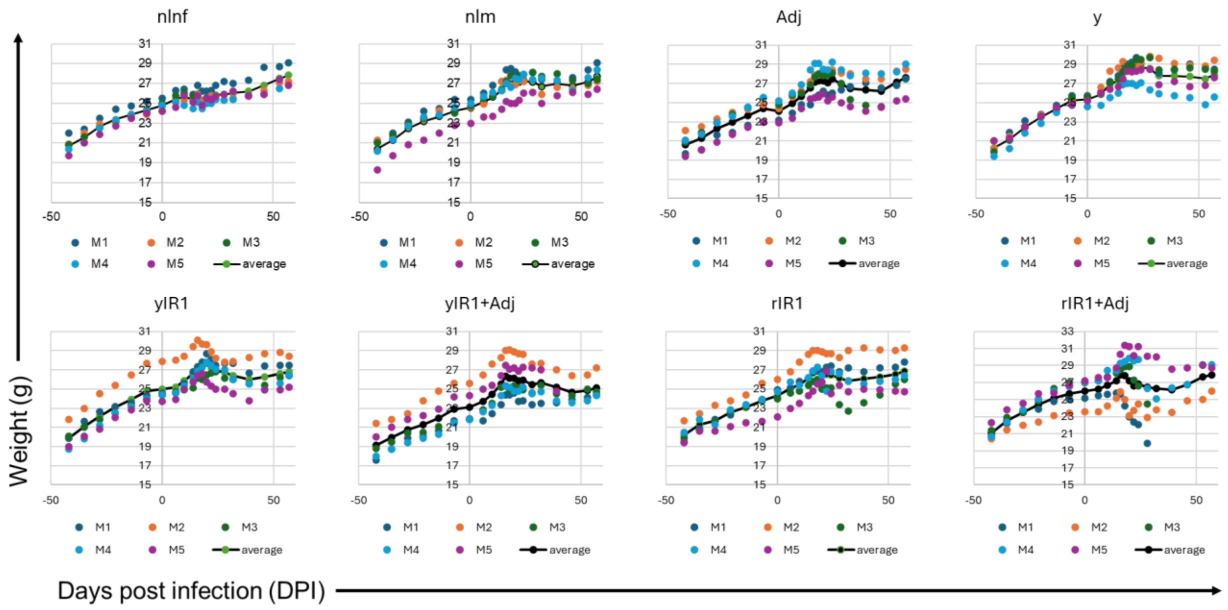


**B**

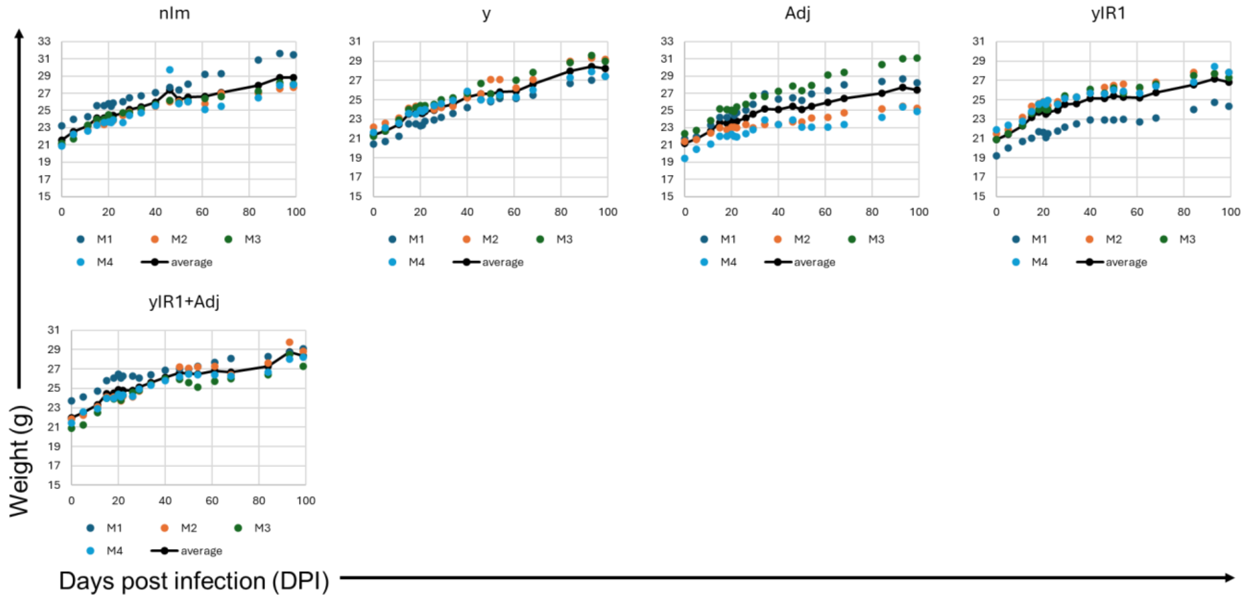


**Figure S6. Mouse weight through vaccination experiments. A)** Mouse weight throughout the prophylactic vaccination experiment. **B)** Mouse weight throughout the therapeutic vaccination experiment. Time 0 corresponds to the time of infection. Weights were recorded every two or three days. M, mouse. nInf, mice non-infected; nIm, mice non-immunized and infected; Adj, mice vaccinate with adjuvant only (IL-12/GM-CSF); y, mice vaccinated with yeast expressing pYD1 empty vector induced with 1% galactose; yIR1, mice vaccinated with yeast expressing pYD1-IR1 induced with 1% galactose; yIR1+Adj, mice vaccinated yIR1 combined with adjuvants; rIR1, mice vaccinated with recombinant IR1 (rIR1); rIR1+Adj, mice vaccinated with rIR1 combine with adjuvants.

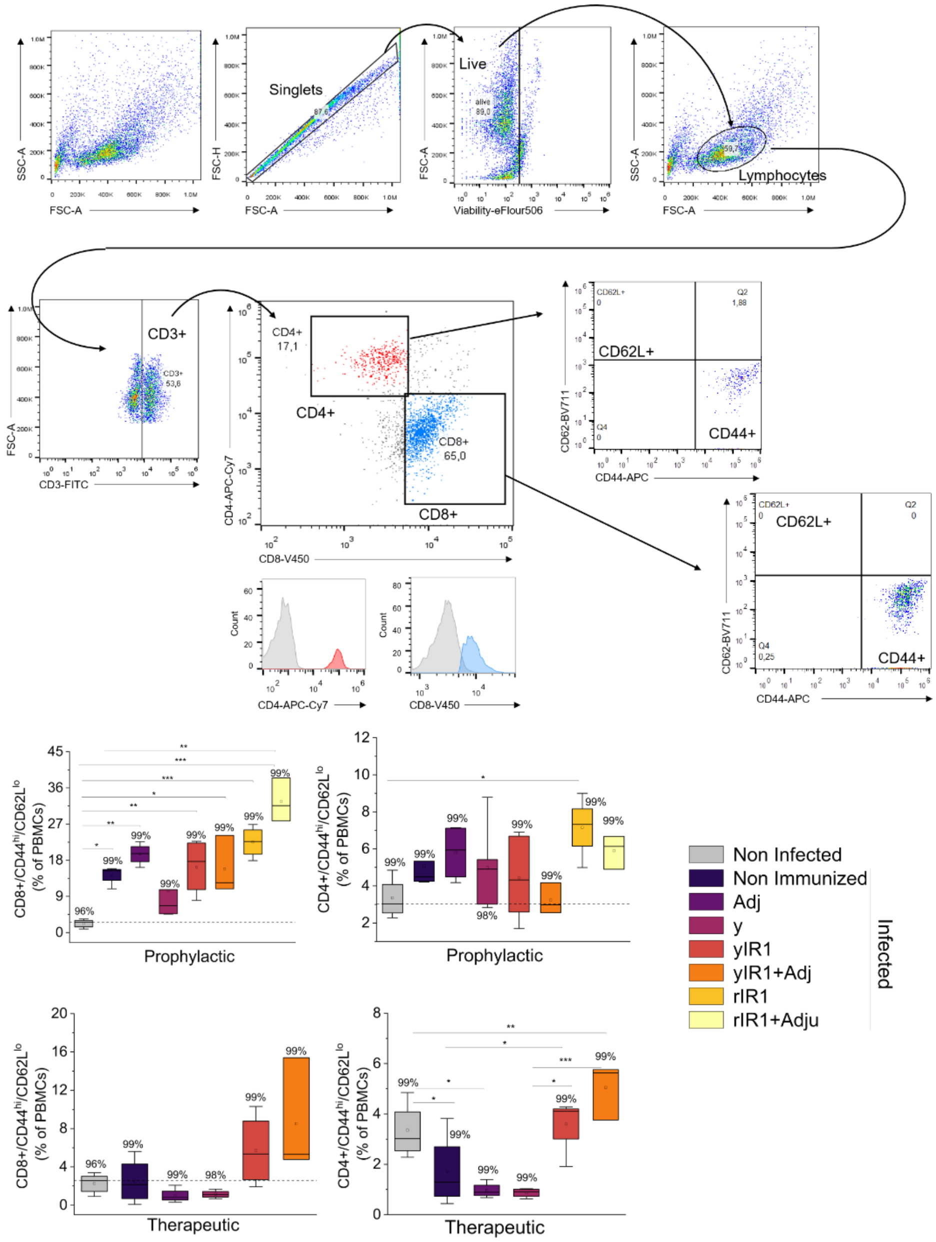


**Figure S7. Gating strategies used for flow cytometry. A)** The gating strategy was performed from PBMCs isolated from yeast-immunized BALB/c mice. Cells were first gated to exclude doublets by selecting singlets based on forward scatter (FSC) area versus height. Subsequently, live cells were identified using viability dye exclusion. From the live-cell population, lymphocytes were gated based on characteristic forward-scatter (FSC) and side-scatter (SSC) properties. T cells were then selected by gating on $CD3^{+}$ events. $CD3^{+}$ cells were then subdivided into $CD4^{+}$ and $CD8^{+}$ T cell populations based on CD4 and CD8 expression, as well as on CD44 and CD62L expression. Histograms show $CD4^{+}$ and $CD8^{+}$ cells compared to unlabelled cells. Numbers shown within each gate indicate the percentage of the parent population. Flow parameters were defined using UltraComp eBeads™ Plus Compensation Beads. **B-C)** Quantification of effector memory CD8+ (CD3+/CD8+/CD44+/CD62L-) or CD4+ (CD3+/CD4+/CD44+/CD62L-) T cells from prophylactic (B) or therapeutic (C) vaccinations.

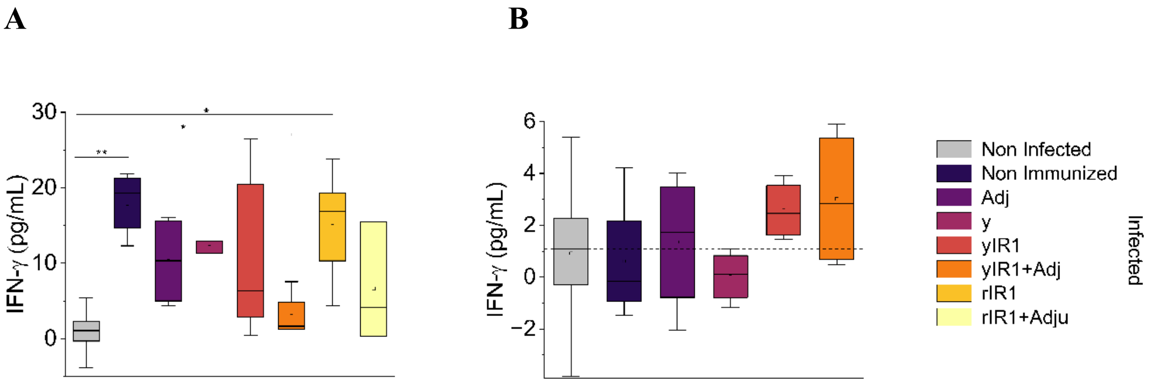


**Figure S8. Quantification of interferon-γ (IFN-γ) in mouse serum by ELISA. A-B)** ELISA quantification of IFN-γ from mouse sera (1:500 dilution) from prophylactic vaccination (A) or therapeutic vaccination (B). The data show the distribution from measurements of 5 mice in plot A and 4 mice in plot B. The horizontal line within the box indicates the median, the box represents the interquartile range (IQR, 25th–75th percentiles), and the whiskers extend to 1.5 times the IQR. The open square within the plot shows the mean. The dotted line shows the median values of non-infected mice. *, $p \leq 0.05$, ** $p \leq 0.01$, *** $p \leq 0.001$. Nonparametric multiple-contrast test with Dunnett-type contrasts was used to compare experimental groups against the non-infected/non-immunized group.

**Table S1. Confirmation of mouse infection by quantification of blood parasitemia.** Parasites were quantified by real-time PCR at 20 days post-infection (dpi) – the first time point at which all mice were detected infected. nInf, mice non-infected; nIm, mice non-immunized and infected; Adj, mice vaccinate with adjuvant only (IL-12/GM-CSF); y, mice vaccinated with yeast expressing pYD1 empty vector induced with 1% galactose; yIR1, mice vaccinated with yeast expressing pYD1-IR1 induced with 1% galactose; yIR1+Adj, mice vaccinated yIR1 combined with adjuvants; rIR1, mice vaccinated with recombinant IR1 (rIR1); rIR1+Adj, mice vaccinated with rIR1 combine with adjuvants. N = 5, i.e., mice per condition. NA, mice not analyzed. dpi, days post-infection. UD, undetected. Values are shown as parasites/mL.

| Day | Mice | nInf | nlm | Adj | y | yIR1 | yIR1 +Adj | rIR1 | rIR1 +Adj |
|---|---|---|---|---|---|---|---|---|---|
| 20 dpi | 1 | UD | 63.85 | 51.84 | 119.52 | 62.08 | 119.39 | 71.02 | 80870.31 |
| | 2 | UD | 68.08 | 85.60 | 80.56 | 143.47 | 73.43 | 55.22 | 2815.70 |
| | 3 | UD | 51.77 | 76.76 | 63.58 | 77.13 | 44.00 | 7.70 | 379.14 |
| | 4 | UD | 102.77 | 77.11 | 53.51 | 74.33 | 76.12 | 32.51 | 178.63 |
| | 5 | UD | 59.01 | 129.61 | 14.85 | 71.56 | 18.93 | 76.58 | 140.91 |

**Table S2. Quantification of blood parasitemia in mice vaccinated with prophylactic IR1 at the endpoint.** Parasites were quantified by real-time PCR at 55 dpi (endpoint). nInf, mice non-infected; nIm, mice non-immunized and infected; Adj, mice vaccinate with adjuvant only (IL-12/GM-CSF); y, mice vaccinated with yeast expressing pYD1 empty vector induced with 1% galactose; yIR1, mice vaccinated with yeast expressing pYD1-IR1 induced with 1% galactose; yIR1+Adj, mice vaccinated yIR1 combined with adjuvants; rIR1, mice vaccinated with recombinant IR1 (rIR1); rIR1+Adj, mice vaccinated with rIR1 combine with adjuvants. N = 5, i.e., five mice per condition. UD, undetected. Background noise of qPCR amplification for blood was 3.4 parasites/mL, intestine tissue was 70.8 parasites/ 10 ng of DNA, and heart was 332.0 parasites/ 10 ng of DNA, as established using the mean of qPCR values of ninf mice. Any amplification above this value is shown, and below it is considered negative (N). NA, mice not analyzed. Values in blood are in parasites/mL, in tissues are parasites/10 ng tissue DNA.

| Tissue | Mice | nInf | nlm | Adj | y | yIR1 | yIR1 +Adj | rIR1 | rIR1 +Adj |
|---|---|---|---|---|---|---|---|---|---|
| Blood 55 dpi | 1 | UD | UD | 98.63 | 204.70 | 167.31 | UD | UD | UD |
| | 2 | UD | 64.61 | 566.32 | 21.86 | 119.30 | 271.33 | UD | 135.55 |
| | 3 | UD | UD | UD | N | N | 61.34 | UD | UD |
| | 4 | UD | 165.73 | 176.407 | UD | 100.33 | 81.46 | N | 182.19 |
| | 5 | UD | N | UD | 293.644 | UD | UD | 170.29 | 115.73 |
| Intestine 55 dpi | 1 | UD | N | 3381.44 | N | N | 176.08 | 674.25 | 265.07 |
| | 2 | UD | N | 42047.30 | 217325.56 | 105.52 | N | 399.01 | 85.99 |
| | 3 | UD | 461.99 | N | N | N | 78.13 | N | 438.94 |
| | 4 | UD | N | 166.19 | 311.40 | 98.46 | NA | 154.22 | NA |
| | 5 | UD | N | NA | N | N | NA | N | NA |
| Heart 55 dpi | 1 | UD | 4158.439 | 605.788 | 669.889 | N | 3149.926 | 413.677 | 13869.065 |
| | 2 | UD | N | 2252.031 | N | N | N | N | 3522.989 |
| | 3 | UD | 532.543 | 1212.458 | 1177.443 | 5743.237 | 1710.438 | 431.961 | 5186.447 |
| | 4 | UD | 6535.153 | N | 992.151 | 6180.891 | 2405.443 | 6779.256 | NA |
| | 5 | UD | 721.887 | NA | 9558.761 | 2426.062 | NA | N | NA |

**Table S3. Confirmation of infection by quantification of blood parasitemia in mice used for therapeutic yIR1 vaccination.** Parasites were quantified by real-time PCR at 5 dpi. nIm, mice non-immunized and infected; Adj, mice vaccinate with adjuvant only (IL-12/GM-CSF); y, mice vaccinated with yeast expressing pYD1 empty vector induced with 1% galactose; yIR1, mice vaccinated with yeast expressing pYD1-IR1 induced with 1% galactose; yIR1+Adj, mice vaccinated yIR1 combined with adjuvants (IL-12/GM-CSF). N = 4, i.e., four mice per condition.

| Group | Sample ID | $C_T$ Mean | parasite/ml | Infected |
|---|---|---|---|---|
| nIm | A1 | 30.130 | 55.10 | Y |
| nIm | A2 | 30.166 | 53.45 | Y |
| nIm | A3 | 30.436 | 42.57 | Y |
| nIm | A4 | 30.084 | 57.29 | Y |
| Adj | B1 | 30.767 | 32.21 | Y |
| Adj | B2 | 28.683 | 186.55 | Y |
| Adj | B3 | 28.564 | 206.12 | Y |
| Adj | B4 | 28.418 | 233.13 | Y |
| y | C1 | 31.146 | 23.42 | Y |
| y | C2 | 29.288 | 112.04 | Y |
| y | C3 | 30.873 | 29.48 | Y |
| y | C4 | 30.762 | 32.36 | Y |
| yIR1 | D1 | 30.613 | 36.68 | Y |
| yIR1 | D2 | 29.137 | 127.16 | Y |
| yIR1 | D3 | 30.111 | 56.01 | Y |
| yIR1 | D4 | 30.717 | 33.60 | Y |
| yIR1+Adj | E1 | 31.182 | 22.72 | Y |
| yIR1+Adj | E2 | 31.470 | 17.82 | Y |
| yIR1+Adj | E3 | 29.881 | 67.97 | Y |
| yIR1+Adj | E4 | 28.516 | 214.59 | Y |

**Table S4. Quantification of parasitemia in mice vaccinated with therapeutic yIR1 in blood and tissues.** Parasites were quantified by real-time PCR at 100 dpi (endpoint). nIm, mice non-immunized and infected; Adj, mice vaccinate with adjuvant only (IL-12/GM-CSF); y, mice vaccinated with yeast expressing pYD1 empty vector induced with 1% galactose; yIR1, mice vaccinated with yeast expressing pYD1-IR1 induced with 1% galactose; yIR1+Adj, mice vaccinated yIR1 combined with adjuvants (IL-12/GM-CSF). N = 4, i.e., four mice per condition. Background noise of qPCR amplification for blood was 3.4 parasites/mL, and for heart was 332.0 parasites/10 ng of DNA, as established using the mean of qPCR values of ninf mice. Any amplification above this value is shown, and below it is considered negative (N). NA, mice not analyzed. Values in blood are parasites/mL; in tissues, parasites/10 ng of tissue DNA.

| Tissue | nIm | Adj | y | yIR1 | yIR1+Adj |
|---|---|---|---|---|---|
| **Blood 100 dpi** | N | 63.55 | 265.64 | 210.49 | 497.65 |
| | N | 152.73 | N | 45.76 | 7.65 |
| | 101.31 | 71.69 | N | 20.39 | 1643.11 |
| | 104.19 | N | 22.15 | 21.55 | 49.32 |
| **Heart 100 dpi** | N | N | N | 3034.42 | 1251.88 |
| | N | N | 12953.42 | N | N |
| | N | N | 624.60 | 563.27 | N |
| | 3678.69 | N | NA | N | 886.13 |

**Table S5. Primers used in this study.** F, forward; R, reverse. Lab ID, Cestari Lab primer database identification number; Lab YID, Cestari Lab yeast database identification number.

| # | Common Name | F/R | Lab ID | Lab YID | Sequence |
|---|---|---|---|---|---|
| 1 | pYD1-BamH1-F | F | 132 | | TTAAGCTTCTGCAGGCTAGTGGTG |
| 2 | pYD1-BamH1-R | R | 133 | | CACTGTTGTTATCAGATCAGCGGG |
| 3 | IR1_pYD1-F | F | 556 | 100 | cgatgacgataaggtaccaggatccATGCTAACTGAAATGAAGCGTG |
| 4 | IR1_pYD1-R | R | 557 | 100 | tgcagaattccaccacactggatccttgaaaatacaaattttcCATGTCTGGTGGGACGTC |
| 5 | IR2_pYD1-F | F | 554 | 99 | cgatgacgataaggtaccaggatccATGACGATGTCGCACGTC |
| 6 | IR2_pYD1-R | R | 555 | 99 | tgcagaattccaccacactggatccttgaaaatacaaattttcCGAAAGAAATGCGGAAAGAG |
| 7 | IR3_pYD1-F | F | 568 | 102 | cgatgacgataaggtaccaggatccATGAAACGCGTGCTGGTG |
| 8 | IR3_pYD1-R | R | 569 | 102 | tgcagaattccaccacactggatccttgaaaatacaaattttcGGTCTGGGTCCAGGTAAA |
| 9 | IR4_pYD1-F | F | 572 | 103 | cgatgacgataaggtaccaggatccATGCGTGCATTTTTGGCC |
| 10 | IR4_pYD1-R | R | 573 | 103 | tgcagaattccaccacactggatccttgaaaatacaaattttcAGTTCCGAGCGAAGTTGCGTA |
| 11 | IR5_pYD1-F | F | 690 | 104 | cgatgacgataaggtaccaggatccATGTTTTACGTTGTTACCTCCTCC |
| 12 | IR5_pYD1-R | R | 691 | 104 | tgcagaattccaccacactggatccttgaaaatacaaattttcTGGAACAAGAATCACAGCGCT |
| 13 | IR1_pET28a-F | F | 707 | | ctggtgccgcgcggcagccatatgATGAAGCGTGTGCTGGTG |
| 14 | IR1_pET28a-R | R | 706 | | agtggtggtggtggtggtgctcgagCTACTCTTCAAGGATGGCGAT |
| 15 | cruzi1 | F | 837 | | ASTCGGCTGATCGTTTTCGA |
| 16 | cruzi2 | R | 838 | | AATTCCTCCAAGCAGCGGATA |
| 17 | cruzi3 | | 839 | | /56-FAM/CACACACTGGACACCAA/3BHQ_1/ |
| 18 | IAC-F | F | 840 | | ACCGTCATGGAACAGCACGTA |
| 19 | IAC-R | R | 841 | | CTCCCGCAACAAACCCTATAAAT |
| 20 | IAC-probe | | 842 | | /5SUN/AGCATCTGTTCTTGAAGGT/3BHQ_1/ |
| 21 | GAPDH-F | F | 863 | | CAATGTGTCCGTCGTGGATCT |
| 22 | GAPDH-R | R | 864 | | GTCCTCAGTGTAGCCCAAGATG |
| 23 | ACTB_F | F | 879 | | GTGACGTTGACATCCG TAAAGA |
| 24 | ACTB_R | R | 880 | | GCCGGACTCATCGTACTCC |
| 25 | IFNg-F | F | 885 | | AGCGGCTGACTGAACTCAGATTGTAG |
| 26 | IFNg-R | R | 886 | | GTCACAGTTTTCAGCTGTATAGGG |
| 27 | Tgfb_F | F | 895 | | TGGAGCAACATGTGGAACTC |
| 28 | Tgfb_R | R | 895 | | GTCAGCAGCCGGTTACCA |
| 29 | IL-4_F | F | 891 | | CGAAGAACACCACAGAGAGTGAGCT |

| | | | | |
|---|---|---|---|---|
| **30** | IL-4_R | R | 892 | GACTCATTCATGGTGCAGCTTATCG |
| **31** | IL-10_F | F | 881 | CTCGTTTGTACCTCTCTCCG |
| **32** | IL-10_R | R | 882 | ATCTCCCTGGTTTCTCTTCC |

**Table S6. Oxford nanopore sequencing statistics.** Sequences with a quality (Q-score) above 7 were used for data analysis.

| Sample | Total number of mapped bases | Total number of alignments | Percentage of mapped reads | Mean sequencing quality (MAPQ) |
|---|---|---|---|---|
| TcYSD 1 | 0.884 Gb | 1842801 | 89.25 | 48.87 |
| TcYSD 2 | 1.345 Gb | 2919913 | 96.14 | 48.58 |
| TcYSD 3 | 1.349 Gb | 2581430 | 96.38 | 49.57 |
| CD_Enrich 1 | 0.482 Gb | 1811657 | 93.61 | 55.04 |
| CD_Enrich 2 | 1.393 Gb | 3184281 | 97.01 | 51.93 |
| CD_Enrich 3 | 0.570 Gb | 2073843 | 94.08 | 53.80 |
| HI_Enrich 1 | 0.438 Gb | 1692527 | 93.40 | 55.65 |
| HI_Enrich 2 | 0.367 Gb | 1603718 | 95.35 | 54.23 |
| HI_Enrich 3 | 0.351 Gb | 1459009 | 92.30 | 55.80 |

**Table S7. Antibodies used in this study.**

| Antibody | Dilution factor | Manufacturer |
|---|---|---|
| Mouse α-Xpress | 1:2000 | Life Technologies |
| rabbit α-Pex14 | 1:5000 | Dr. Marilyn Parsons, Seattle Childrens |
| rabbit α-glycosome (PGKA,B,C) | 1:5000 | Dr. Marilyn Parsons, Seattle Childrens |
| goat α-mouse-HRP | 1:5000 | BioRad |
| donkey α-rabbit | 1:5000 | BioRad |
| mouse α-His-tag | 1:2000 | Invitrogen |
| FITC Hamster Anti-Mouse CD3e | 1:400 | BD Biosciences |
| APC-Cy™7 Rat Anti-Mouse CD4 | 1:400 | BD Biosciences |
| V450 Rat anti-Mouse CD8a | 1:400 | BD Biosciences |
| APC Rat Anti-Mouse CD44 | 1:400 | BD Biosciences |
| BV711 Rat Anti-Mouse CD62L | 1:400 | BD Biosciences |
| Goat anti-Human IgG (H+L) Cross-Adsorbed Secondary Antibody, Alexa Fluor Plus 405 | 1:1000 | Thermo Fisher Scientific |

**Description of Data files S1 to S4.**

**Data file S1. YSD antigen-antibody screen enriched genes.** The dataset shows genes enriched for binding to IgG antibodies from CD patients, HI, or both. FC, fold-change; CPM, counts per million.

**Data file S2. IR expressed by *T. cruzi* infectious forms.** The dataset shows IRs with evidence for expression in metacyclic trypomastigotes (MT), amastigotes (AM), or cell-derived trypomastigotes (CT). Expression was obtained via tandem mass tag labelling of *T. cruzi* proteins, followed by mass spectrometry-based identification. IR_ID, IR identification, which combines gene ID with the IR genome starting position.

**Data file S3. IRs with epitopes identified in the EpAtlas and IEDB.** The dataset lists IRs whose amino acid sequences match epitopes identified in EpAtlas, along with the percentage of patient sera that react with each epitope and the number of IRs each epitope matches. It also shows the IRs with sequences matching epitopes identified in the IEDB. The amino acid sequence of each epitope match is also shown.

**Data file S4. List of all IRs and data used for vaccine target prioritization.** The dataset lists all 1,047 identified IRs, along with their nucleotide and amino acid sequences, their presence in discrete typing units (DTUs) I to VI, their abundance in each *T. cruzi* infectious form as identified by tandem mass tag labelling of *T. cruzi* proteins followed by mass spectrometry-based identification, and their functional annotation. The YSD antigen-antibody screen enrichment is shown with log2 fold-change (log2FC), log2 counts per million (log2CPM), and Benjamini-Hochberg (BH) adjusted *p*-value. The data indicate which proteins are human homologs, members of multigene families (MGFs), and those expressed in all *T. cruzi* infectious forms. The 143 IRs prioritized as vaccine targets are also indicated.
